# Iowa Brain-Behavior Modeling Toolkit: An Open-Source MATLAB Tool for Inferential and Predictive Modeling of Imaging-Behavior and Lesion-Deficit Relationships

**DOI:** 10.1101/2024.07.31.606046

**Authors:** Joseph C. Griffis, Joel Bruss, Stein F. Acker, Carrie Shea, Daniel Tranel, Aaron D. Boes

**Affiliations:** Department of Pediatrics, Carver College of Medicine, University of Iowa, 200 Hawkins Drive, Iowa City, IA, 52242, USA; Department of Neurology, Carver College of Medicine, University of Iowa, 200 Hawkins Drive, Iowa City, IA, 52242, USA; Medical Scientist Training Program, Carver College of Medicine, University of Iowa, 200 Hawkins Drive, Iowa City, IA, 52242, USA; Department of Psychological and Brain Sciences, University of Iowa, 200 Hawkins Drive, Iowa City, IA, 52242, USA; Department of Psychiatry, Carver College of Medicine, University of Iowa, 200 Hawkins Drive, Iowa City, IA, 52242, USA; Iowa Neuroscience Institute, University of Iowa, 200 Hawkins Drive, Iowa City, IA, 52242, USA

## Abstract

The traditional analytical framework taken by neuroimaging studies in general, and lesion-behavior studies in particular, has been inferential in nature and has focused on identifying and interpreting statistically significant effects within the sample under study. While this framework is well-suited for hypothesis testing approaches, achieving the modern goal of precision medicine requires a different framework that is predictive in nature and that focuses on maximizing the predictive power of models and evaluating their ability to generalize beyond the data that were used to train them. However, few tools exist to support the development and evaluation of predictive models in the context of neuroimaging or lesion-behavior research, creating an obstacle to the widespread adoption of predictive modeling approaches in the field. Further, existing tools for lesion-behavior analysis are often unable to accommodate categorical outcome variables and often impose restrictions on the predictor data. Researchers therefore often must use different software packages and analytical approaches depending on whether they are addressing a classification vs. regression problem and on whether their predictor data correspond to binary lesion images, continuous lesion-network images, connectivity matrices, or other data modalities. To address these limitations, we have developed a MATLAB software toolkit that supports both inferential and predictive modeling frameworks, accommodates both classification and regression problems, and does not impose restrictions on the modality of the predictor data. The toolkit features both a graphical user interface and scripting interface, includes implementations of multiple mass-univariate, multivariate, and machine learning models, features built-in and customizable routines for hyper-parameter optimization, cross-validation, model stacking, and significance testing, and automatically generates text-based descriptions of key methodological details and modeling results to improve reproducibility and minimize errors in the reporting of methods and results. Here, we provide an overview and discussion of the toolkit’s features and demonstrate its functionality by applying it to the question of how expressive and receptive language impairments relate to lesion location, structural disconnection, and functional network disruption in a large sample of patients with left hemispheric brain lesions. We find that impairments in expressive vs. receptive language are most strongly associated with left lateral prefrontal and left posterior temporal/parietal damage, respectively. We also find that impairments in expressive vs. receptive language are associated with partially overlapping patterns of fronto-temporal structural disconnection, and that the associated functional networks are also similar. Importantly, we find that lesion location and lesion-derived network measures are highly predictive of both types of impairment, with predictions from models trained on these measures explaining ∼30-40% of the variance on average when applied to data from patients not used to train the models. We have made the toolkit publicly available, and we have included a comprehensive set of tutorial notebooks to support new users in applying the toolkit in their studies.

**Key Points:**

1. We describe a new MATLAB toolkit for inferential and predictive modeling of imaging datasets.
2. We demonstrate the application of the toolkit to real lesion, network, and behavioral datasets.
3. We have made the toolkit publicly available to the research community.

## 1. Introduction

Lesion-symptom mapping is an important methodology in neurology and cognitive neuroscience (Damasio and Damasio, 1989; Rorden and Karnath, 2004). While early approaches to lesion-symptom mapping relied on post-mortem examinations of brain tissue to establish associations between lesion location and neurological deficits, modern approaches rely primarily on 3-dimensional lesion images derived from non-invasive neuroimaging technologies such as computerized tomography (CT) and magnetic resonance imaging (MRI). The statistical methods used to map the observed neurological deficits to the underlying neuroanatomy can be divided into two primary categories: 1) mass univariate statistical approaches that separately model the lesion-symptom relationships for each predictor (e.g., voxel), and (2) multivariate statistical approaches that model the lesion-symptom relationship for all predictors (e.g., voxels) simultaneously within a single unified model (Moore et al., 2024). Traditional approaches to modeling lesion-symptom relationships are *inferential* in nature – models are fit to a single dataset, and inferences about the underlying lesion-symptom relationships are drawn based on the results of statistical tests on the model parameters. While inferential modeling is critical for drawing evidence-based conclusions about how observed symptoms relate to the underlying neuroanatomy, robust predictive models are necessary to achieve the goal of translating basic lesion research into clinical applications (Poldrack et al., 2020). There is accordingly growing interest in developing and evaluating *predictive* models that learn generalizable lesion-symptom relationships and that allow for prediction of cognitive and behavioral outcomes in independent datasets. The development of research tools that facilitate the development, evaluation, and application of such models is therefore of high importance.

However, currently available tools for lesion-symptom research are designed primarily for inferential modeling applications and offer limited built-in functionality for predictive applications (DeMarco et al., 2018; Pustina et al., 2017a; Zhang et al., 2014). While there is also widespread interest in incorporating lesion-derived network features such as parcel and tract-level summary statistics, disconnection matrices, and functional connectivity networks into lesion-symptom models (Bowren et al., 2022; Griffis et al., 2019, 2021; Olafson et al., 2023; Sperber et al., 2022), existing tools are often inflexible in terms of the types of inputs that they can accommodate (e.g., binary NIFTI images, specific parcellations, etc.). The lack of publicly available research tools that feature a wide range of modeling approaches, customizable predictive modeling options, and the ability to handle a diverse range of input features is particularly detrimental given both the growing emphasis on developing generalizable predictive models of lesion-symptom relationships, and the increasingly common incorporation of lesion-derived features, such as lesion-derived networks inferred from ‘connectome’ data, or disconnection matrices into lesion-symptom modeling workflows.

Here, we describe a new publicly available software tool designed to support both inferential and predictive modeling of brain-behavior relationships, with an emphasis on applications in lesion research. This tool includes multiple multivariate and mass-univariate modeling approaches appropriate for modeling both continuous (i.e., regression approaches) and categorical (i.e., classification approaches) outcomes, features built-in statistical testing frameworks to evaluate statistical significance at the model-level and coefficient-level, and includes built-in and customizable routines for predictive modeling that include hyper-parameter optimization, repeat nested cross-validation with repartitioning, cross-validation permutation testing, and the ability to combine the outputs of different models to improve predictive performance (i.e., model stacking). Importantly, this tool can flexibly accommodate different types of predictor data, including binary lesion segmentations, continuous lesion-network maps, anatomical summaries of the lesion (e.g., tract disconnection and parcel damage estimates), and adjacency matrices (e.g., “disconnectomes”), allowing for identical analysis approaches to be applied across a range of predictor modalities and facilitating the rigorous and systematic comparison of models utilizing different types of input features. Finally, this tool features both a simple-but-flexible graphical user interface (GUI) and an easily customizable scripting interface, making it maximally accessible while also providing a high level of customizability, flexibility, and scalability.

## 2. Tool Overview

### Software Requirements

The tool is written in the MATLAB programming language (TheMathWorks), and it was developed and tested using MATLAB version R2022b. The tool requires the MATLAB Statistics and Machine Learning Toolbox for core model implementations (required for core toolbox functionality), the MATLAB Image Processing Toolbox for NIFTI file input/output (required for reading/writing NIFTI image files), the MATLAB Bioinformatics Toolbox (required for False Discovery Rate Correction), and the MATLAB Parallel Processing Toolbox for parallel processing functionality (not required but recommended). The GUI was created using the MATLAB App Designer, and tutorial scripts for the scripting interface are provided as MATLAB Live Notebooks. The toolkit is designed to leverage the MATLAB Parallel Processing Toolbox whenever possible, greatly reducing the time required to run computationally intensive analyses such as hyper-parameter optimization, permutation testing, or bootstrap resampling, although parallel processing is not required.

### Data Modalities

Behavioral outcomes of interest may be either continuously distributed (e.g. a score on a neuro-psychological test) or categorical (e.g. the presence vs. absence of a deficit). Accordingly, this tool features comprehensive modeling options appropriate for both continuous and binary categorical behavioral outcomes (see Modeling Approaches section).

The predictor data in traditional lesion-symptom mapping analyses correspond to binary voxel-based lesion segmentations. However, newer methods such as “lesion-network mapping” and “disconnectome-mapping” often produce continuously varying estimates of network (dis)connectivity (Joutsa et al., 2022a; Sperber et al., 2022). There is therefore a need for comprehensive modeling tools that can accommodate diverse predictor modalities.

Accordingly, the toolkit does not impose restrictions on the modality of the predictor data. Through the GUI, it includes dedicated functions to load and format imaging data contained in subject-level NIFTI files, connectivity/adjacency matrices stored in text or CSV files, and arbitrary predictor arrays stored in text or CSV files. Through the scripting interface, it can accommodate any arbitrary data matrix where rows correspond to observations and columns correspond to predictors.

### Modeling Approaches

The toolkit provides users with an array of different modeling options, allowing them to select the most appropriate modeling approach given their research goals and constraints. This also allows for direct comparisons of different modeling approaches within the same software package, facilitating methodological research. Importantly, it allows users to straightforwardly evaluate the stability of results across multiple modeling approaches, as recommended by recent guidelines for conducting lesion-symptom research (Moore et al., 2024).

#### Model Implementations

The toolkit includes the following mass-univariate modeling approaches: Pearson correlations (implemented using the MATLAB *corr()* function), *t*-tests (equal or unequal variances; implemented using the MATLAB *ttest2()* function), Brunner-Munzel tests (implemented using scripts hosted here: https://github.com/robisoe/Brunner-Munzel-test-for-matlab), ordinary least squares regressions and binary logistic regressions (implemented using the MATLAB *fitglm()* function), and proportional subtraction analyses.

The toolkit also includes implementations of the following multivariate modeling approaches: ridge-and LASSO-penalized regressions implemented using the MATLAB *fitrlinear()* function, ridge and LASSO penalized linear classifiers implemented using the MATLAB *fitclinear()* function, partial least squares regression and partial least squares classification implemented using the MATLAB *plsregress()* function, linear and non-linear support vector regressions and classifiers implemented using the MATLAB *fitrsvm()* and *fitcsvm()* functions, and ensemble regression and classification models (e.g., bagged or boosted tree models) implemented using the MATLAB *fitrensemble()* and *fitcensemble()* functions.

#### Cross-validation and Hyper-parameter Optimization Routines

The toolkit includes built-in cross-validation routines for evaluating out-of-sample prediction performance. The default approach is *k*-fold cross-validation, where the dataset is partitioned into a specific number of groups represented by the variable *k,* and each fold is iteratively used as a “test” set while the remaining folds are used to train the model (e.g. a 5-fold partition of 100 subjects would have 20 subjects per fold, with 80 used for training and 20 used for testing at each iteration). Leave-one-out (LOO) cross-validation, where the model is iteratively fit on data from *N*-1 “train” samples and the held-out sample is used as the “test” set, is also supported. While both approaches provide estimates of out-of-sample prediction performance, the *k*-fold approach is often faster due to needing to fit only *k* models instead of *N* models. The two approaches also differ with respect to the bias vs. variance of the estimates, with *k*-fold being generally recommended for predictive analyses (Poldrack et al., 2020; Scheinost et al., 2019), motivating our decision to use it by default. For *k*-fold approaches, repeated *k-* fold splits with repartitioning of train/test folds may be performed to obtain estimates of performance across different partitions of the data in order to mitigate partition noise. These routines allow researchers to evaluate how well predictive models can be expected to generalize to data not used to train the models.

Hyper-parameters in multivariate models are variables that control model behavior and that must be pre-specified before training the model. For PLS approaches, hyper-parameters correspond to the *number of PLS components* used to fit the model. For SVMs, they correspond to the *BoxConstraint*, *KernelScale*, and (for SVR) *epsilon* parameters (see MATLAB *fitrsvm*()/*fitcsvm*() documentation). For regularized linear models, they correspond to the regularization parameter *lambda* that controls the shrinkage of the model coefficients. For bagged tree ensemble models, they correspond to the *NumLearningCycles* and *MinLeafSize* parameters that control the number of weak learners and minimum number of leaf node observations, respectively (see MATLAB *fitrensemble*()*/fitcensemble*() documentation). Since hyper-parameters are not learned during model training, cross-validation provides a principled way to tune the hyper-parameters of multivariate models by identifying the hyper-parameter value(s) that minimize out-of-sample prediction error.

Hyper-parameter tuning can be performed within a cross-validation analysis by using a nested cross-validation approach (Scheinost et al., 2019). Nested cross-validation involves first partitioning the full dataset into “outer train” and “outer test” sets, as in typical cross-validation schemes. Then, the “outer train” set is further partitioned into “inner train” and “inner test” sets, with these “inner” partitions being used to optimize model hyperparameters (i.e., by identifying the hyperparameters that minimize prediction errors on the “inner test” set) and the “outer” partitions being used to fit and evaluate the performance of models trained with optimized hyperparameters (e.g., for a nested 5-fold approach with 100 subjects, the 80 subjects in the “outer training” set at a given iteration would be split into “inner training” and “inner test” sets with 16 subjects per fold, with 64 used for training and 16 used for testing at each iteration). For cross-validation analyses with multivariate models, nested cross-validation is used by default.

For all cross-validation analyses, user-specified data transformation steps such as variable standardization and confound regression (see next sub-section) are performed within the cross-validation loop to prevent data leakage and ensure separation of training and test datasets (Poldrack et al., 2020; Scheinost et al., 2019). Optionally, the user can stratify cross-validation partitions according to a user-specified stratification variable (e.g., the outcome variable) to ensure representation of different portions of the data distribution across partitions (Varoquaux et al., 2017). For continuous stratification variables, stratification is performed by dividing the dataset into groups based on the quartiles of the stratification variable, while for categorical stratification variables, stratification is performed according to the corresponding categories. Stratification may be particularly useful for highly skewed behavioral data or categorical variables with heavily imbalanced classes.

#### Confound Mitigation Strategies

Researchers often wish to remove variance associated with one or more nuisance variables, such as lesion volume or age, from the outcome variable prior to performing inferential lesion-symptom mapping analyses (Moore et al., 2024). Accordingly, the toolkit includes a flexible confound regression option that allows researchers to specify confounds that they want to regress from the outcome prior to proceeding with the modeling analyses. If selected, any confound variables specified by the researcher as predictors will be regressed from the outcome prior to performing the main analyses, and the residuals from this regression will be designated as the new outcome variable. For mass-univariate analyses using general linear models (i.e., ordinary linear regression, logistic regression), confound variables are not regressed from the outcome, but are instead included as covariates in the model. We note that for cross-validation analyses, all confound regressions are performed on the “training” data, and the resulting models are then applied to the “test” data to obtain the residualized outcome in order to avoid data leakage.

For lesion-symptom mapping analyses using voxel-level lesion data as predictors, the direct total lesion volume control (DTLVC) approach described by Zhang and colleagues (2014), which transforms each patient’s lesion by dividing the voxel values by the square root of the lesion volume such that the transformed lesion array has a unit norm equal to 1, may also be used. This approach is intended to downweigh the contributions of large lesions at each voxel and is less conservative than the nuisance regression approach to controlling lesion volume effects in support vector regression models (DeMarco et al., 2018). However, we note that this approach was originally developed for support vector models, and its effects on other models have not been thoroughly evaluated. An important avenue for future work is therefore to systematically characterize the effects of data processing choices such as DTLVC on different modeling strategies. We hope to facilitate these types of methodological studies by making this toolkit available to the research community.

### Statistical Inference

#### Mass-Univariate Models

Mass-univariate analyses typically rely on null hypothesis significance testing (NHST) to identify imaging features (e.g., voxels, edges, etc.) that exhibit statistically significant bivariate associations with the outcome of interest (Mirman et al., 2017). For mass-univariate analyses, classical parametric significance tests and non-parametric permutation-based significance tests are both supported by the toolkit, with non-parametric permutation-based tests being utilized by default.

Permutation testing involves constructing an empirical null distribution for the statistic of interest by iteratively shuffling the outcome variable to break any true relationship between the predictors and outcome before refitting the model to the shuffled outcomes. Statistical significance is determined by comparing the statistic obtained from the original analysis against the null distribution obtained from the permutation analyses. Specifically, the *p*-value is computed as the number of null statistics at least as extreme as the observed statistic, divided by the number of permutation iterations plus one.

Permutation testing for mass-univariate analyses can be performed either at the level of each individual model (e.g., voxel-wise permutation tests) or across all models simultaneously (e.g., whole-brain permutation tests). The toolkit performs whole-brain permutation tests using the continuous family-wise error rate (cFWER) control method (Mirman et al., 2017) by default. This approach, which is an extension of the commonly used “Max-T” approach, controls the probability that a specified number of results are false positives at the nominal alpha level across all models simultaneously (e.g., 100 voxels at *a*=0.05). It was chosen as the default because it has been shown to outperform similar methods such as cluster extent correction or false discovery rate (FDR) correction in mass-univariate analyses of small patient samples (Mirman et al., 2017). It is applied at voxel count thresholds of [1, 10, 50, 100, 500, 1000], providing the user with results across a range of thresholds as recommended by Mirman and colleagues (2017).

Individual model-level (e.g., voxel-level) permutation *p*-values can also be obtained, although this is disabled by default due to the large number of permutation iterations required to achieve predictor-level *p*-values small enough for FWER corrections to be viable in voxel-based analyses. Additionally, classical parametric individual model-level *p*-values can also be obtained for mass-univariate analyses. For either permutation-derived or parametric model-level *p*-values, multiple comparisons correction can be applied via either the Benjamini-Hochberg method for false discovery rate (FDR) control (Benjamini and Hochberg, 1995) or the Bonferroni-Holm method for FWER control (Aickin and Gensler, 1996).

#### Model-level Tests for Multivariate Models

Currently, there is not consensus on how to best perform statistical inference in multivariate lesion-symptom mapping analyses, and a wide range of approaches are used in the literature (Corbetta et al., 2015; DeMarco et al., 2018; Pustina et al., 2017a; Zhang et al., 2014). We take the perspective that in the context of multivariate models, there are two “levels” of statistical inference that are relevant to model interpretation. The first is “model-level” inference, which focuses on the interpretability of the full model (e.g., “does the model capture a meaningful relationship between the predictors and the outcome?”). The second is “coefficient-level” inference, which focuses on the interpretability of the individual model coefficients (e.g., “which voxels show reliable contributions to the model?”). While many multivariate lesion-symptom mapping studies do not evaluate or report model-level tests, we believe that model-level tests are nonetheless important for contextualizing coefficient-level results – the interpretation of a significant model coefficient should be contingent on the quality of the model fit.

For this reason, we take the perspective that explicit model-level statistical tests are needed to contextualize the interpretation of coefficient-level results in inferential applications of multivariate models. We propose a two-step workflow for lesion-symptom inference on inferential (i.e., “explanatory”) multivariate models. This approach involves first performing model-level evaluations to inform model interpretation (including determining whether the model should be interpreted at all), and then proceeding to perform coefficient-level inference if the model passes at least one model-level evaluation. Model-level evaluations also allow for inferences about the joint effects of all predictors simultaneously (e.g., at the level of the full voxel coefficient map – Pustina et al., 2017a) even if the individual model coefficients do not survive multiple comparisons corrections and prevent inferences about the effect of any individual predictor. Model-level evaluations can be performed in the toolkit using either permutation testing, or full-sample cross-validation testing.

Permutation tests are performed on the in-sample model fit and intend to determine whether the model fits better than expected if there were truly no relationship between the predictors and the outcome variable. This is achieved by first generating a distribution of model fit statistics obtained from empirical null models and then comparing that distribution against the observed model fit statistic. This is performed in the same way as described for mass-univariate analyses in the previous section (i.e., the outcome is shuffled to break the true association with the predictors), with the exception that a single model is fit using all predictors at each permutation iteration. For regression models, the model mean squared error (MSE) is used as the model fit statistic, while for classification models, the model area under the ROC curve is used (Poldrack et al., 2020). A significant *p-*value indicates that the model fits the data better than expected if there were no true relationship between the predictor matrix and the outcome variable, i.e. that the model is capturing a potentially meaningful relationship between the predictors and the outcome, justifying further inference on model coefficients.

Full-sample cross-validation tests can be used to determine whether predictions generated by the model for new data points (i.e., data not used to train the model) exhibit a statistically significant relationship with the true observed outcomes at the level of the full sample, and is the approach used to test for map-level significance in the LESYMAP software package (Pustina et al., 2017a). This approach aims to avoid caveats of in-sample model fit statistics (e.g., overfitting) in high-dimensional multivariate models by using a nested cross-validation framework to optimize model hyper-parameters and obtain out-of-sample predictions for each observation in the dataset. For each repetition of the *k*-fold cross-validation procedure, one out-of-sample prediction is obtained for each observation in the full dataset. If repeated cross-validations are performed, then the predicted outcomes are summarized across repetitions of the cross-validation procedure by taking the mean or median (i.e., for regression) or the mode (i.e., for classification) across all repetitions, resulting in a single summary out-of-fold prediction for each observation in the full dataset. The group-level association between these out-of-fold predictions and the true observed outcomes is then evaluated using either Pearson correlation (i.e., for continuous outcomes) or Fisher’s exact test (i.e., for categorical outcomes) to obtain a parametric *p*-value reflecting the probability of obtaining an association at least as extreme as the observed association if there were no true relationship between the model predictions and the observed outcomes. A significant *p-*value indicates that the model is capturing a statistically meaningful association between the predictor data and the outcome variable across all out-of-fold predictions for the dataset, justifying further inference on the model coefficients. We note that while this test employs a cross-validation framework, the result of this test should not be used as the primary measure of a model’s out-of-sample prediction performance since it is computed using the out-of-fold predictions for the whole dataset (Poldrack et al., 2020). In addition, for regression models, the cross-validation correlation does not consider the magnitudes of prediction errors but only the linearity of the relationship between the predicted and observed outcomes (Poldrack et al., 2020; Scheinost et al., 2019). The full-sample cross-validation results should therefore be regarded as a group-level summary estimate of the association between model predictions and observed outcomes across the full sample under study, reflecting the ability of the model to capture meaningful associations that generalize beyond the data it was trained on (Pustina et al., 2017a).

While both of these approaches utilize a null hypothesis significance testing framework, they nonetheless take different perspectives on model-level inference – permutation tests concern the *in-sample* quality of model fit, while full-sample cross-validation tests concern the expected *out-of-sample* generalizability of the model, which are different but complementary perspectives on model inference (Bzdok and Yeo, 2017). In-sample model fit statistics (e.g., R-squared, classification accuracy) obtained from multivariate models should be interpreted with caution due to the tendency of these models to overfit to the training sample, especially for highly flexible models such as support vector models using non-linear kernels (Poldrack et al., 2020). Given that such models can produce good fits even in the absence of a true relationship (e.g., see permutation test results using in-sample MSEs for the non-linear support vector models in **Supplementary Figures 7-9**), we consider the test of *out-of-sample* generalizability to be most informative since it explicitly evaluates whether the model is capturing a relationship that generalizes beyond the training dataset. However, the full-sample cross-validation test approach may not be viable in smaller datasets, and permutation testing has the advantage of testing the exact model that is being used for coefficient-level inference (i.e., in cross-validation testing, many models are trained and evaluated, which may differ from the final group-level model). We therefore recommend that researchers thoughtfully consider whether one approach (or both approaches) is most appropriate given their dataset, research question, modeling strategy, and other factors.

#### Coefficient-level Tests for Multivariate Models

The second level of inference for multivariate models is coefficient-level inference, which seeks to draw conclusions about the effects of individual predictors in the model. For example, after obtaining a statistically significant result from a model-level test, a researcher may wish to draw inferences about the effects of individual predictors in the model (e.g., individual voxel peaks). The statistical significance of the individual predictors (e.g., voxels) is then evaluated separately from the statistical significance of the overall model.

Existing tools use varying approaches to coefficient-level inference that include voxel-level permutation tests (Zhang et al., 2014), cluster-level permutation tests (DeMarco et al., 2018), or avoiding it altogether in favor of model-level inferences on a sparse model (Pustina et al., 2017a). These approaches have potential drawbacks. For example, in high-dimensional analyses such as voxel-based lesion analyses, voxel-level permutation tests require a huge number of permutation iterations before it is possible to obtain *p*-values small enough to survive multiple comparisons corrections at typical alpha levels (e.g., 0.05) -for a typical analysis including 156,919 2mm isotropic voxels, more than 3 million permutation iterations would be required before it was possible to obtain *p*-values small enough to pass Bonferroni correction at the *p <* 0.05 threshold. On the other hand, cluster-level permutation tests require that the user specify an *a priori* uncorrected *p*-value threshold for cluster formation (discussed in detail in Mirman et al., 2017), and they only allow for the inference that there is an effect for at least one of the voxels within each significant cluster (Woo et al., 2014). While other approaches, such as the permutation-based “Max-T” approach, have been applied to multivariate lesion-symptom mapping analyses in the literature (Ivanova et al., 2021a; Jiang and Gong, 2024), it’s unclear if/when these and related approaches such as the cFWER approach (Mirman et al., 2017) are appropriate for multivariate models where all individual predictor effects (1) are potentially of interest, (2) are conditional on the values of all other predictors in the model and don’t directly reflect the strength of each predictor’s bivariate association with the outcome, and (3) do not directly reflect the strength of the relationship captured by the full model (e.g., a null model that performs much worse than the observed model could nonetheless have larger model coefficients at some voxels, which can’t happen in the mass-univariate context). Accordingly, while both voxel-level permutation tests and cFWER permutation tests are implemented in the toolkit and can be used with multivariate models, we caution against their use for the reasons outlined above, and they are disabled by default.

Resampling techniques such as the jack-knife and bootstrap (Griffis et al., 2019; Kohoutová et al., 2020; Yourganov et al., 2018) can be used to estimate the stability of the model coefficients across repeated re-fitting to different data samples, and they provide a straightforward way to identify statistically robust predictors in multivariate models. This is our preferred approach for supporting coefficient-level inference in multivariate models. Unlike permutation tests, which aim to determine whether the *magnitudes* of the model coefficients are greater than expected if there were no relationship between the predictor matrix and outcome variables, resampling approaches aim to estimate whether the *population parameters* for the model coefficients are reliably different from zero. These approaches naturally accommodate the inter-dependence among model coefficients in multivariate models and do not depend on the assumption of “voxel-level exchangeability” required for permutation tests. Accordingly, the toolkit uses bootstrap testing as the default approach to coefficient-level inference for multivariate models. Bootstrap testing, which uses random resampling with replacement to iteratively refit the model and estimate the distribution of model parameters, allows for confidence interval estimation (parametric or non-parametric) on model coefficients, and for parametric significance testing using *z*-statistics computed from the mean and standard deviation (i.e., bootstrap standard error) of the bootstrap distribution (Kohoutová et al., 2020). The Benjamini-Hochberg FDR procedure and the Bonferroni-Holm FWER procedure are used to correct for multiple comparisons, and results are thresholded using both procedures by default to provide both liberal and conservative corrected *p*-value estimates on model coefficients.

#### Evaluating Prediction Performance for Multivariate Models

The evaluation of model performance in an independent dataset is necessary to establish the predictive power of a model, and is an important first step towards translating a model into real-world applications (Poldrack et al., 2020; Scheinost et al., 2019). However, it is often not possible or practical for researchers to obtain an independent dataset that is well-suited for evaluating the out-of-sample prediction performance of their model. Cross-validation strategies provide a way for researchers to evaluate the predictive performance of their model(s) within a single dataset. As described in the earlier section on cross-validation, these approaches involve partitioning the dataset into non-overlapping “train” and “test” sets, fitting the model to the “train” set, and then evaluating the out-of-sample prediction performance by applying the fitted model to the “test” set.

Multiple estimates of out-of-sample predictive performance are provided for predictive analyses using cross-validation. For regression models, these include the mean squared error of prediction, explained variance score, sum-of-squares-based prediction *R*^2^, and Pearson correlation between predicted and observed values (Poldrack et al., 2020). For classification models, these include the overall classification accuracy, classification accuracy for each class, and the area under the ROC curve (Poldrack et al., 2020). Under the default repeat *k*-fold nested cross-validation scheme, these measures are computed for each test fold for each repeat of the cross-validation procedure. These measures, along with the individual test set predictions, model coefficients, and hyper-parameters for each test fold for each repeat are stored along with the full cross-validation partitions in a file that is saved by default at the end of the full cross-validation analysis.

Permutation testing can be used to evaluate the statistical significance of cross-validation performance in predictive analyses (Scheinost et al., 2019). For permutation testing, the outcome variable is permuted prior to each permutation iteration, which corresponds to a full run of the entire cross-validation procedure (e.g., if the original analysis used 5 repeats of 5-fold cross-validation, then a single permutation iteration would correspond to 5 repeats of 5-fold cross-validation with a permuted outcome variable). For each permutation iteration, performance measures are summarized across folds and across repeats to generate a single summary performance measure for the full cross-validation run (e.g., average MSE over all folds and repeats), yielding an *N_permutations* x 1 distribution of summary performance measures which are then compared against the corresponding summary performance measure from the original cross-validation analysis to yield a permutation *p*-value. For regression approaches, the performance measure is the MSE, while for classification approaches, the performance measure is the area under the ROC curve (Poldrack et al., 2020). A significant *p-*value indicates that average out-of-sample prediction performance of the model reliably outperforms those of empirical null models, supporting conclusions about the predictive power of the model.

### Additional Features for Predictive Modeling

#### Model Stacking

The toolkit also allows for users to fit multiple different models (e.g., lesion-symptom mapping, lesion-network mapping, disconnection matrix analyses, etc.) to predict the same outcome in a given sample, and then combine the predictions from these models with the goal of improving prediction performance (Olafson et al., 2023; Pustina et al., 2017b). The simplest approach involves summarizing (e.g., averaging) predictions across different models (Olafson et al., 2023), while more complex approaches involve “stacking” the predictions across models into an *N_subjects*-by-*N_models* predictor matrix that is then used as input to a second “meta-model” that is trained (using the original CV partitions) to predict outcomes using the stacked predictions from the base models as predictors (Pustina et al., 2017a). The implementation in the toolkit allows the user to run each individual “base” model with a “model stacking” flag, generating identical cross-validation partitions for different models applied to the same dataset (e.g., lesion, disconnection, etc.). Alternately, the cross-validation partitions for a previously trained model can be designated for training subsequent models on the same patient sample. After running the individual “base” models, the entire cross-validation routine can be re-run using the predictions obtained from the base models as predictor features for the new “meta-model”.

#### Applying Trained Models to New Datasets

While cross-validation techniques such as *k-*fold and leave-one-out cross-validation enable the evaluation of model generalizability within a single dataset, these techniques are nonetheless limited by the characteristics of the data sample under study. When practical, it is therefore ideal to apply the fully trained models from one dataset to observations obtained from a fully independent dataset to evaluate model performance.

Within the toolkit, the application of trained models to new datasets facilitated by a built-in prediction function that simply takes as inputs the result output file containing the fully trained model from the original run (these are saved together by default) along with a new predictor matrix with the same columns as the original predictor matrix used to train the model (e.g., voxels extracted using the same brain mask), and that outputs predictions for each observation in the new dataset. If any data transformations were applied to the predictor data during the original analysis (e.g., variable standardization, DTLVC, etc.), then these transformations are automatically applied to the new data using the parameters obtained from the training dataset in the original analysis.

### Toolkit Interface

The toolkit features a simple, yet flexible graphic user interface (GUI) that can be called from the MATLAB command window. This allows users to interactively select input data, configure analysis parameters, define the output location, and run the analysis without needing to write any code. A fully configured analysis using the main GUI is shown in **Figure 1**. An overview of the options implemented in the GUI along with an example workflow are provided in the User Manual that is included with the toolkit. Analysis steps are printed to the command window, and after analyses are complete, main modeling results are printed to the command window and are also saved to a text file along with the standard result outputs in a pre-specified result directory. There are two separate GUIs, one for running individual modeling analyses, and the other for running stacked modeling analyses.

**Figure 1.**
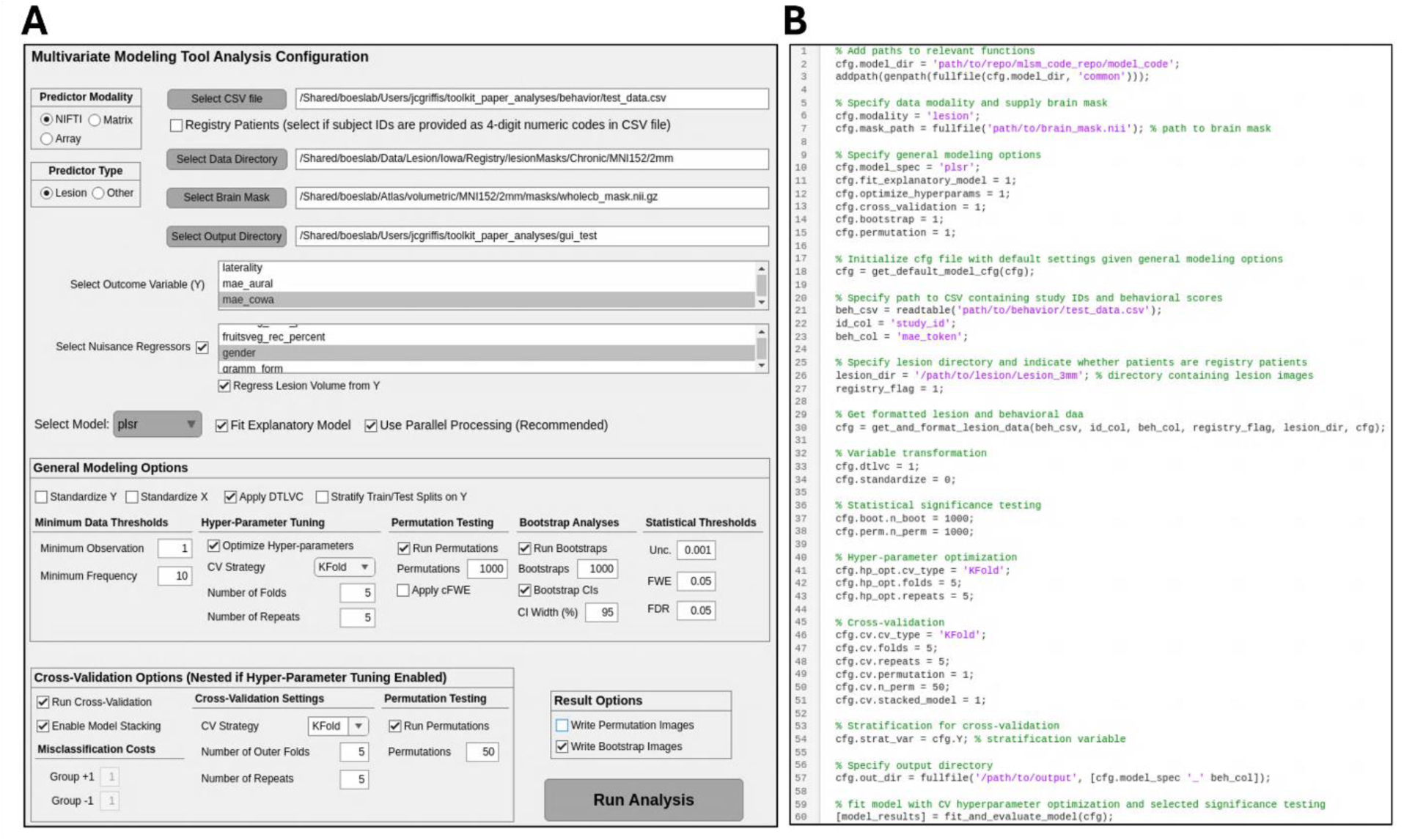
Main Toolkit Interfaces. **A.** A complete analysis configured using the GUI. **B.** The same analysis configured using the scripting interface. Both panels show a fully configured lesion-deficit modeling analysis that includes hyper-parameter optimization, repeat nested 5-fold cross-validation with permutation testing, and fitting a final inferential model to the full dataset with permutation testing for model-level significance and bootstrap testing for coefficient-level significance. The GUI also shows selection of nuisance regressors for illustration purposes. Using either interface, it is simple to quickly configure and run complex modeling analyses. We note that use of the scripting interface is not required, and the code snippet is provided simply to illustrate what it looks like to specify a complete end-to-end analysis using the scripting interface.

The toolkit also features a scripting interface that allows for users to flexibly design and implement custom analyses by defining and modifying fields and corresponding values within a MATLAB structure array (struct) type variable called “cfg” (short for configuration). Once specified, this *“*cfg” variable can be provided as the input argument to core toolkit functions to run custom analyses with user-defined options and parameters. This facilitates the creation of custom analysis scripts and simple wrapper functions that can run analyses at scale via e.g., high-performance computing clusters. An example end-to-end script configuring and running an analysis that includes hyper-parameter optimization, repeat nested 5-fold cross-validation with permutation testing, and fitting a final inferential model to the full dataset with permutation testing for model-level significance and bootstrap testing for coefficient-level significance is shown in **Figure 1B**. As noted earlier, multiple MATLAB Live Notebooks are included with the toolkit. These provide step-by-step tutorials that illustrate how to implement custom inferential and predictive analyses using different modeling implementations and predictor modalities (i.e., lesion data formatted as NIFTI images, functional lesion-network maps formatted as NIFTI images, parcel-wise disconnection matrices formatted as text files, ROI summary measures formatted as text files) via the scripting interface, and provide a detailed overview of the result outputs.

### Toolkit Output

#### Results files

Analysis results are saved as .mat files containing a “model_results” struct for inferential/full-sample modeling analyses (along with the trained model for multivariate models), and a “cv_results” struct for cross-validation analyses. These files contain performance measures, relevant analysis parameters, and other relevant information for evaluating and interpreting the model results. All initial analysis parameters (including the random seed and state) and relevant data (e.g., model hyper-parameters) are saved with the final modeling results by default. This ensures that all information needed to reproduce the results is available even if the script that produced the results is modified or lost after running the analysis. Users can also opt to save the full outputs of permutation testing and resampling procedures (i.e., these were used to generate the histograms in **Figures 4-7**), but they are not saved by default due to the large size of the resulting files.

#### Imaging files

For NIfTI predictor inputs, different NIfTI images can be output along with the results files depending on the specific analysis performed. These include unthresholded and statistically thresholded (e.g., FDR or FWE-corrected) model coefficient and test statistic maps. The maps will be output in the same space and resolution as the original input data and accompanying user-specified brain mask. For lesion analyses, a lesion frequency map (i.e., a heatmap indicating the frequency of damage at each voxel) is produced by default along with a map encoding the correlation between lesion status at each voxel and overall lesion volume (see **Figure 2**). These maps provide valuable information about the spatial distribution of lesions and about the relationship between lesion topography and lesion size in the sample under study, and they can provide important context for understanding potential confounds and for interpreting results. For adjacency matrix inputs that are provided along with a table of node coordinates, model coefficients can be output as .node and .edge files for viewing in external viewers (e.g., ball-and-stick images in **Figure 3B,E**).

**Figure 2.**
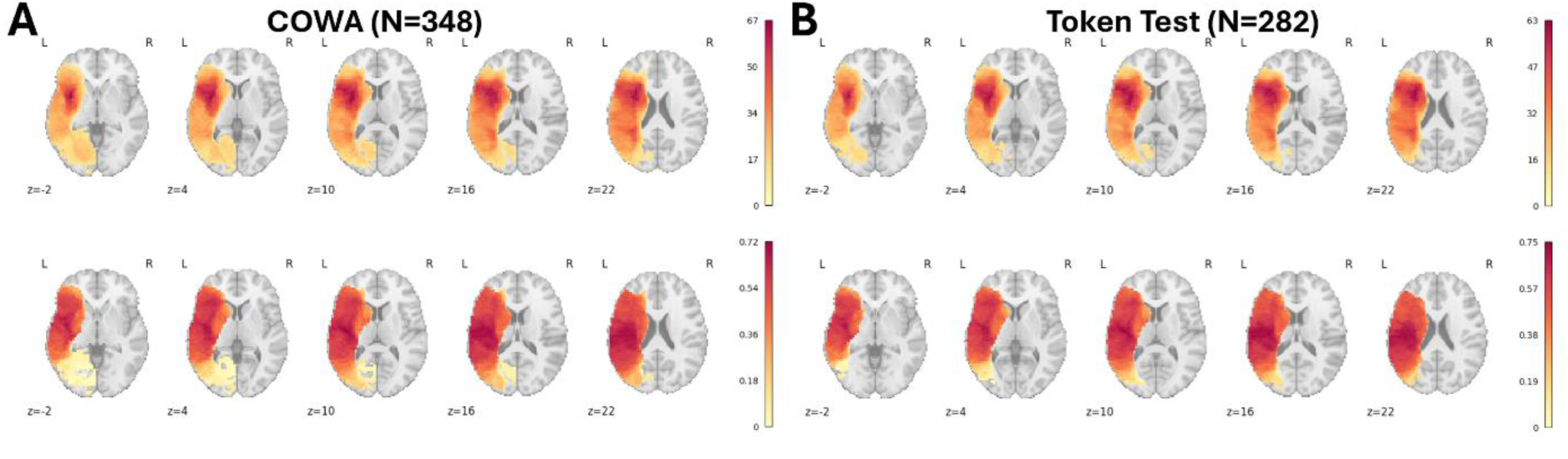
Lesion frequency maps and lesion volume correlation maps. **A.** The top row shows lesion frequencies for the sample of patients with data for COWA (i.e., voxels shown in hot colors are damaged more frequently). The bottom row shows correlations between voxel lesion statuses and lesion volume (i.e., damage to voxels shown in hot colors is associated with larger lesions). **B.** The same maps are shown for patients with data for the Token Test.

**Figure 3.**
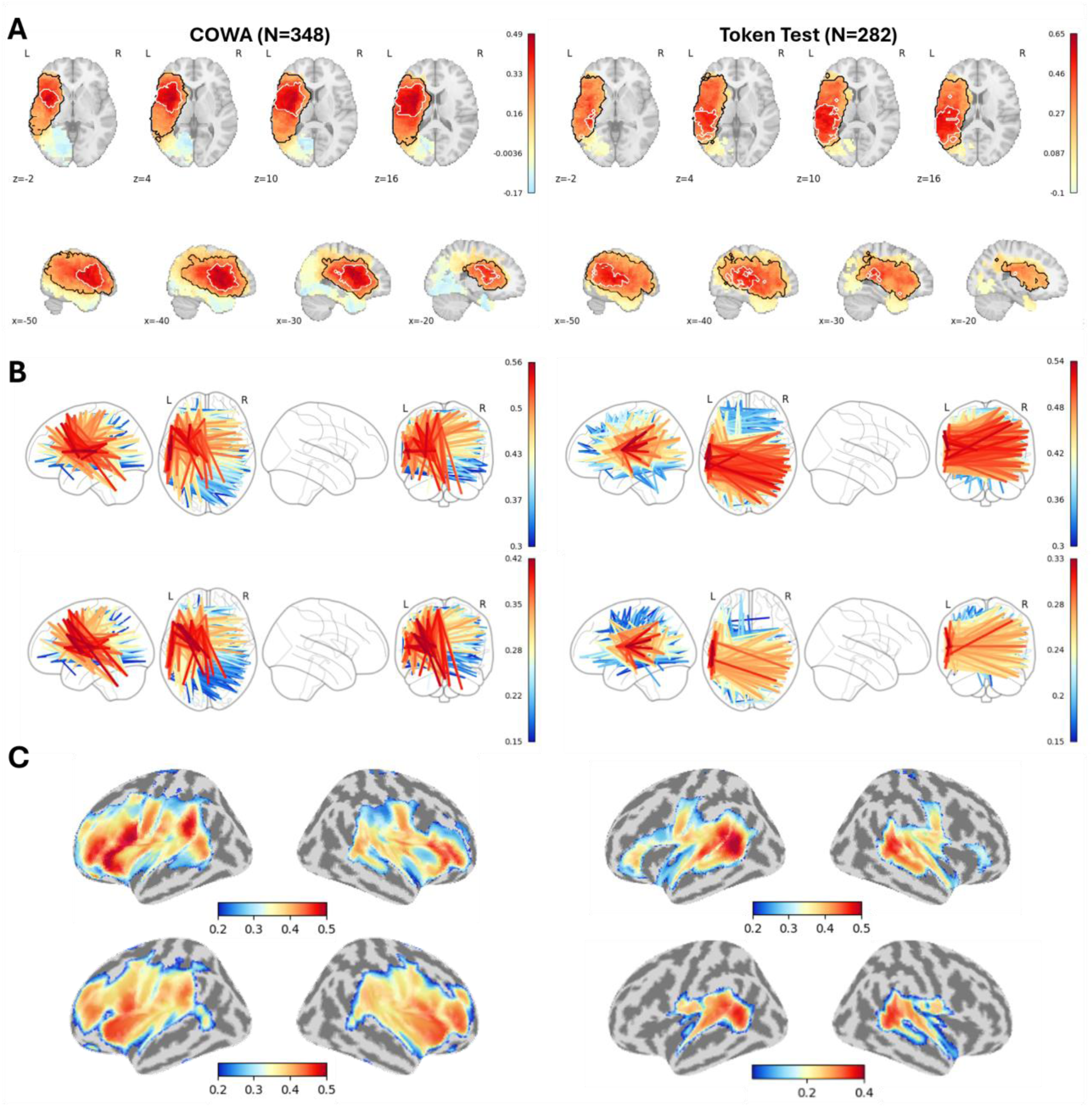
Mass-univariate lesion, structural disconnection, and functional lesion-network analyses of expressive and receptive language impairments. **A.** Results of analyses using voxel-based lesion maps. The maps show unthresholded correlations between voxel lesion statuses and COWA scores. Voxels surviving the FWE*p*<0.05, *v*=100 threshold for the analyses without lesion volume regression are outlined in black, and voxels surviving this threshold for the analyses with lesion volume regression are outlined in white. **B**. Results of analyses using parcel-to-parcel structural disconnection matrices. Top rows show results for analyses without lesion volume regression, and bottom rows show results of analyses with lesion volume regression. **C.** Results of analyses using fLNM maps. Top rows show results for analyses without lesion volume regression, and bottom rows show results of analyses with lesion volume regression. All analyses are thresholded using the FWE*p*<0.05, *v*=100 threshold.

#### Automated Report Generation

The toolkit also automatically generates “boiler plate” text in paragraph form that describes the main details of the methods and results to facilitate accurate reporting and enhance reproducibility. These text descriptions are stored as fields in the “model_results” struct and are also saved as text files in the specified results directory. Examples are provided in **Supplementary Material 1** for several of the analyses reported later in the paper.

### Limitations

There are several limitations to the current version of the toolkit that should be noted. The toolkit does not currently feature the cluster-level multiple comparisons correction for voxel-level permutation tests that is featured in the SVR-LSM Toolkit (DeMarco et al., 2018). Nonetheless, it does feature the option for whole-brain cFWER correction of permutation test results, providing an alternative to voxel-level multiple comparisons corrections, although this is primarily intended for mass-univariate models (Mirman et al., 2017).

For classification models, only binomial outcomes are supported in the initial release, although support for multinomial outcomes could be added in future releases.

While ensemble models are currently supported by the toolkit, they are currently restricted to predictive modeling applications and do not perform statistical testing on model fits or model coefficients. However, they do generate estimates of predictor importance using the MATLAB functions *oobPermutedPredictorImportance()* for “Bag” models and *predictorImportance()* for boosted ensemble models, allowing for the characterization of important predictors.

Finally, the toolkit currently is only available in the MATLAB programming language, which requires a paid license to use, and depends on several MATLAB toolboxes. Even so, MATLAB is very commonly used in neuroimaging and lesion-symptom research, and several other existing toolkits are implemented in MATLAB as well (e.g., NiiStat, SVR-LSM). In the future, the toolkit could potentially be adapted to other languages such as Python or R to increase accessibility.

### Comparisons with Other Tools

To show how the functionality of the IBB Toolkit relates to that of existing software tools, we constructed a table summarizing the functionality of the IBB Toolkit and three other toolkits designed for the statistical analysis of lesion data and/or imaging data obtained from patients with brain lesions (Table 1). We note that this table is far from comprehensive, as there are often nuanced differences among the specific methods, options, result outputs, and other features/constraints of the toolkits described in the table. Nonetheless, it is apparent from the table that compared to existing tools, the IBB Toolkit offers greater flexibility in terms of input data modalities, a more diverse array of modeling options, greater support for predictive modeling analyses, and support for both graphical and scripting interfaces.

**Table 1.** Summary of toolkits for the analysis of lesion and imaging datasets.

| Software | Mass-Univariate Analyses | Multivariate Regression | Multivariate Classification | Hyperparameter Tuning | Predictive Modeling | Predictor Modalities | User Interfaces | Language | Dependencies |
| --- | --- | --- | --- | --- | --- | --- | --- | --- | --- |
| LESYMAP | Brunner-Munzel Test, T-Test, Linear Regression, Chi-Square Test | SCCAN, SVR | None | SCCAN | SCCAN (Limited Within-Sample Cross-Validation, Cross-Dataset Prediction) | Binary Lesion Masks | Scripting | R | ANTSR |
| SVR-LSM Toolkit | None | SVR | None | SVR | None | Binary Lesion Masks | GUI | MATLAB | SPM12, libsvm, MATLAB Statistics and Machine Learning Toolbox, MATLAB Parallel Processing Toolbox |
| NiiStat | GLM | SVR | SVM | SVR, SVM | SVR, SVM (Limited Within-Sample Cross-Validation) | Binary Lesion Masks, Cerebral Blood Flow (ASL), Resting-State (fMRI), Task (fMRI), Diffusion (MRI) | GUI | MATLAB | SPM12, libsvm, Microsoft Excel |
| IBB Toolkit | Pearson Correlation, Brunner-Munzel Test, T-Test, Linear Regression, Logistic Regression, Proportional Subtraction | PLS, Ridge, LASSO, SVR, RF | PLS, Ridge, LASSO, SVM, RF | All Multivariate Models | All Multivariate Models (Within-Sample Cross-Validation, Cross-Dataset Prediction) | Any observation-by-predictor matrix; includes functionality for input/output of NIFTI images and .node/.edge files | Scripting, GUI | MATLAB | MATLAB Statistics and Machine Learning Toolbox, MATLAB Image Processing Toolbox, MATLAB Bioinformatics Toolbox, MATLAB Parallel Processing Toolbox |

## 3. Example Analyses

### Datasets

To illustrate the functionality of the toolkit, we performed both inferential and predictive modeling analyses using real lesion and behavioral data from a sample of patients with left hemispheric brain lesions drawn from the Iowa Lesion Registry. Two behavioral measures from the Multilingual Aphasia Battery were used -the Controlled Oral Word Association Test (COWA) and the Token Test. The COWA is a commonly used measure of expressive language function, while the Token Test is a commonly used measure of receptive language function. To demonstrate toolkit functionality, we employed both regression and classification approaches. For classification models, previously published cut-off scores corresponding to impairment thresholds (Rey et al., 1999) were used to define impaired vs. not impaired group labels. This allowed for the evaluation of both regression and classification approaches using the same dataset. A total of 348 patients had data for the COWA, and a total of 282 patients had data for the Token Test. Dataset demographics are summarized in Table 2.

**Table 2.** Demographic data. For Age (years), Education (years), Handedness, and TSO (time since onset - years), data are shown as mean (SD). For etiology fields, data are shown as counts. Stroke (I) = Ischemic Stroke, Stroke (H) = Hemorrhagic Stroke. Lesion volume is shown as voxel counts for original resolution (1mm^3^) voxels.

| Language Measure | Age | % Male | % White | Education | Handedness | TSO | Stroke (I) | Stroke (H) | Tumor | Other | Lesion Volume |
| --- | --- | --- | --- | --- | --- | --- | --- | --- | --- | --- | --- |
| COWA | 50.88 (15.61) | 54 | 95 | 12.61 (4.21) | 81.22 (52.47) | 3.10 (5.68) | 219 | 57 | 30 | 42 | 39585.48 (54922.28) |
| Token Test | 51.15 (14.4) | 62 | 95 | 12.73 (3.99) | 79.50 (55.0) | 3.04 (3.21) | 183 | 46 | 19 | 34 | 42577.83 (58379.34) |

Lesion masks were segmented and registered to the MNI-152 brain template as described in previous publications using data from the Iowa Lesion Registry (Bowren et al., 2022). Due to the large number of computationally intensive modeling analyses performed in this study, lesion data were resampled to 3mm isotropic resolution for analyses of voxel-based lesion images to reduce computational and storage space requirements. We note that downsampling is not required for lesion analyses in the toolkit, but it greatly reduces computational time and storage space requirements, and in our experience, it does not meaningfully change the results (i.e., results for the main analyses reported in the paper were nearly identical when performed using 1mm isotropic resolution lesion data). We also note that many common analysis techniques reduce the inferential resolution below that of the lesion data by imposing cluster-size thresholds (e.g., DeMarco and Turkeltaub, 2018), smoothing the data during modeling (e.g., Pustina et al., 2018), or summarizing the lesion data in terms of overlaps with pre-defined structures (e.g., Griffis et al., 2021), and the application of a low-pass spatial filter to the lesion data (i.e., smoothing) may actually improve performance in lesion-symptom mapping analyses (Ivanova et al., 2021). Accordingly, we do not believe that it is problematic to utilize lightly downsampled lesion data for analyses unless there is reason to believe that the effects of interest require single millimeter spatial resolution to be detectable.

The Lesion Quantification Toolkit (Griffis et al., 2021) was used to generate parcel-level disconnection matrices for each patient by embedding each patient’s lesion mask into the HCP-1065 streamline tractography atlas (Yeh, 2022) and identifying streamlines that intersected the lesion using the DSI_Studio software package as described in (Griffis et al., 2021). The parcellation was constructed by combining a 200-region functional parcellation of the cortex (Schaefer et al., 2018) with subcortical parcels from the Freesurfer DTK subcortical parcellation, cerebellar parcels from the Cerebellum-SUIT parcellation (Diedrichsen, 2006), and the brainstem parcel from the Harvard-Oxford anatomical atlas (https://fsl.fmrib.ox.ac.uk/fsl/fslwiki/Atlases). Functional lesion-network maps (Joutsa et al., 2022a) were generated by using each patient’s lesion as a seed region in resting-state functional connectivity analyses conducted in the GSP1000 normative subject sample described in previous studies (Joutsa et al., 2022b) using the principal component functional disconnection method (Pini et al., 2021), yielding for each patient a single continuously varying network map reflecting the strength of each voxel’s normative functional connectivity to the lesion.

#### Lesion Frequency and Lesion Volume Correlation Maps

Lesion frequency maps and lesion volume correlation maps were automatically generated for the lesion analyses. Maps for the sample of patients with data for the COWA are shown in **Figure 2A**. Analogous maps are shown for the sample of patients with data for the Token Test in **Figure 2B**. The peak lesion overlaps for both patient samples were located in the left lateral prefrontal cortex, underlying white matter, and basal ganglia (**Figure 2**, top rows). The peak correlations between voxel lesion status and lesion volume for both patient samples were located along the posterior portion of the sylvian fissure and included ventral somato-motor regions, primary auditory regions, and posterior insular regions (**Figure 2B**, bottom row).

#### Analysis 1: Mass-Univariate Lesion-Behavior Mapping

First, we ran a traditional mass-univariate lesion-symptom mapping analysis using the “municorr” option in the toolkit. This approach computes point-biserial linear correlations between the lesion status at each voxel and the outcome measure of interest. We set the “minimum lesion affection” threshold to 10 (i.e., a voxel must be lesioned in at least 10 patients to be included in the analysis), and we performed the analyses both with and without lesion volume regression to illustrate the effects of lesion volume regression on the resulting statistical maps. We also performed mass-univariate correlations between each language outcome and (1) the parcel-to-parcel structural disconnection matrices, and (2) the fLNM maps. For all analyses, statistical significance was determined using the cFWER permutation testing method (Mirman et al., 2017) using 10,000 permutation iterations. Disconnection and fLNM analyses were also performed with and without lesion volume regression. Results were considered significant if they survived an FWE-corrected *p-*value threshold of *p*<0.05 and a voxel count threshold of *v*=100.

Results are shown in **Figure 3**. Analyses were run on a Linux system with 16 cores and 251GB of memory. The lesion analysis for COWA completed in 0.51 minutes, and the lesion analysis for Token Test completed in 0.39 minutes. Without lesion volume regression, both lesion-behavior analyses yielded large swaths of significant voxels throughout the left MCA territory, although the strongest correlations for COWA were located more anteriorly in lateral prefrontal regions and the underlying deep white matter, (**Figure 3A**) while the strongest correlations for the Token Test were located more posteriorly in temporal/parietal regions and the underlying deep white matter (**Figure 3A**). Lesion volume exhibited significant correlations with both the COWA (*r*=0.30, *p*<0.001) and Token Test (*r*=0.52, *p*<0.001). To mitigate lesion volume effects, we regressed lesion volume from each outcome and performed the mass-univariate analyses using the residualized outcomes. After lesion volume regression, the significant voxels for COWA vs. Token Test more clearly localized to frontal vs. temporal/parietal structures that included the deep white matter, respectively (**Figure 3A**). Notably, analyses of both COWA and Token Test implicated voxels in canonical “language” areas (Friederici and Gierhan, 2013) and in regions that have previously been identified as “white matter bottlenecks” in the deep prefrontal and deep temporal white matter (Griffis et al., 2017; Mirman et al., 2015; Turken and Dronkers, 2011).

Analyses using the structural disconnection matrices revealed that impairments on the COWA and Token Test were associated with widespread patterns of structural disconnection involving frontal, temporal, parietal, and sub-cortical/cerebellar regions (**Figure 3B**). Lesion volume regression had minimal impact on either disconnection pattern (**Figure 3B** – top rows vs. bottom rows), and the thresholded upper triangular portions of the matrices from analyses with vs. without lesion volume regression were highly similar (COWA: Dice Coefficient = 0.73; Token Test: Dice coefficient = 0.61). The disconnection patterns associated with impairments on COWA and Token Test were also highly similar to each other (Dice coefficient = 0.91) and remained similar with lesion volume regression (Dice coefficient = 0.59).

Analyses using the fLNM maps revealed that impairments on the COWA and Token Test were associated with lesions exhibiting normative functional connectivity to similar fronto-temporo-parietal networks (**Figure 3C**; Dice coefficient = 0.63). Lesion volume regression had minimal impact the COWA fLNM pattern (Dice coefficient = 0.85) but substantially reduced the extent of the Token Test fLNM pattern (**Figure 3B** – top rows vs. bottom rows; Dice coefficient = 0.42) such that it only included areas proximal to the lesion sites identified in the lesion analyses with lesion volume regression (**Figure 3A**).

Results for lesion-behavior analyses using other mass-univariate methods included in the toolkit (i.e., Brunner-Munzel Test, Unequal Variance T-Tests, Logistic Regressions) are provided in **Supplemental Figure 1**. Lesion-behavior results were highly consistent across all mass-univariate methods. Together, the mass-univariate results suggest that impairments on the COWA and Token Test localize to distinct lesion locations and are associated with partially overlapping patterns of structural and functional disconnection. They also demonstrate the utility of lesion volume regression for increasing the specificity of localizations in the presence of correlations between lesion volume and the behavioral outcomes of interest.

#### Analysis 2: Multivariate Lesion-Behavior Regression

Next, we performed multivariate regression analyses using partial least squares (PLS) regression models using the “plsr” option in the toolkit. We opted to perform both inferential and predictive analyses in order to both (a) identify lesion locations with statistically significant model coefficients, and (b) to evaluate the out-of-sample predictive power of models trained on lesion location. To mitigate the effects of lesion volume without removing variance from the outcome variables, the lesion data were transformed using the direct total lesion volume control (DTLVC) approach (Zhang et al., 2014).

The inferential modeling approach involved fitting the PLS models to the full dataset, performing model-level tests, and then performing coefficient-level tests to identify significant lesion predictors. Model-level tests were performed by evaluating the cross-validation correlation across the full sample using the average out-of-fold predictions from 5 repeats of a nested 5-fold cross-validation analysis, and by evaluating the in-sample model fit relative to empirical null models using permutation tests with 1000 permutation iterations. The number of PLS components to include in each model was determined using 5 repeats of 5-fold cross-validation; components were added to the model until the addition of a new component increased the average out-of-sample MSE (Abdi, 2010; Griffis et al., 2019).

All multivariate analyses were run using the UI Argon high-performance computing cluster. The full multivariate lesion analysis for COWA completed in 30.45 minutes, and the full multivariate lesion analysis for Token Test completed in 24.62 minutes. We note that runtime varied substantially depending on the modeling approach used (e.g., the RBF kernel SVR lesion analysis reported in **Supplemental Figure 3** completed in 847.15 minutes).

Results are shown in **Figure 4**. Model-level tests using the full sample cross-validation correlation (**Figure 4A** - scatterplots) and permutation tests (**Figure 4A** – histograms) yielded statistically significant results. We proceeded to evaluate the model coefficients for the full inferential model using bootstrap *z-*tests (Kohoutová et al., 2020) with 1000 bootstrap iterations. Overall, the coefficient maps produced by the inferential modeling analyses (**Figure 4B**) were highly consistent with the results obtained from the mass-univariate correlation analyses with lesion volume regression (**Figure 3A**), supporting the conclusion that the strongest lesion predictors of impairments on the COWA vs. Token Test exhibit distinct frontal vs. temporal localizations, although we note that the lesion predictors identified by the multivariate analyses using DTLVC extended beyond the more restricted localizations identified by the mass-univariate analyses with lesion volume regression (**Figure 3A**).

Next, we performed predictive analyses to evaluate the ability of the PLSR models to generalize beyond the samples used to train them. These analyses involved performing 5 repeats of nested 5-fold cross-validation, with hyper-parameter tuning being performed in the inner loop and evaluating the average out-of-sample prediction performance (i.e., averaged across folds and repeats). To determine the statistical significance of the average cross-validation performance estimates, a permutation test with 50 permutation iterations was performed on the full cross-validation procedure. The predictive analyses indicated that both models explained ∼35-40% of the variance on average in unseen patient samples (**Figure 4C** – bar graphs), which was statistically significant as indicated by the permutation tests of the full cross-validation procedures (**Figure 4C** – histograms). Similar prediction performance was observed for other types of multivariate models, and the use of DTLVC generally improved prediction performance for most models evaluated relative to not using DTLVC (**Supplementary Figures 2-3, 6-7**). This is consistent with the results reported by Zhang and colleagues (2014), but more work is necessary to fully characterize the effects of DTLVC on predictive performance and localization accuracy for different types of multivariate models. Together, these results provide insights into the ability of models trained on voxel-based lesion location data to predict chronic impairments on these tests in unseen patient samples and indicate that models trained on lesion location can account for nearly 40% of the performance variation when applied to unseen patient samples.

**Figure 4.**
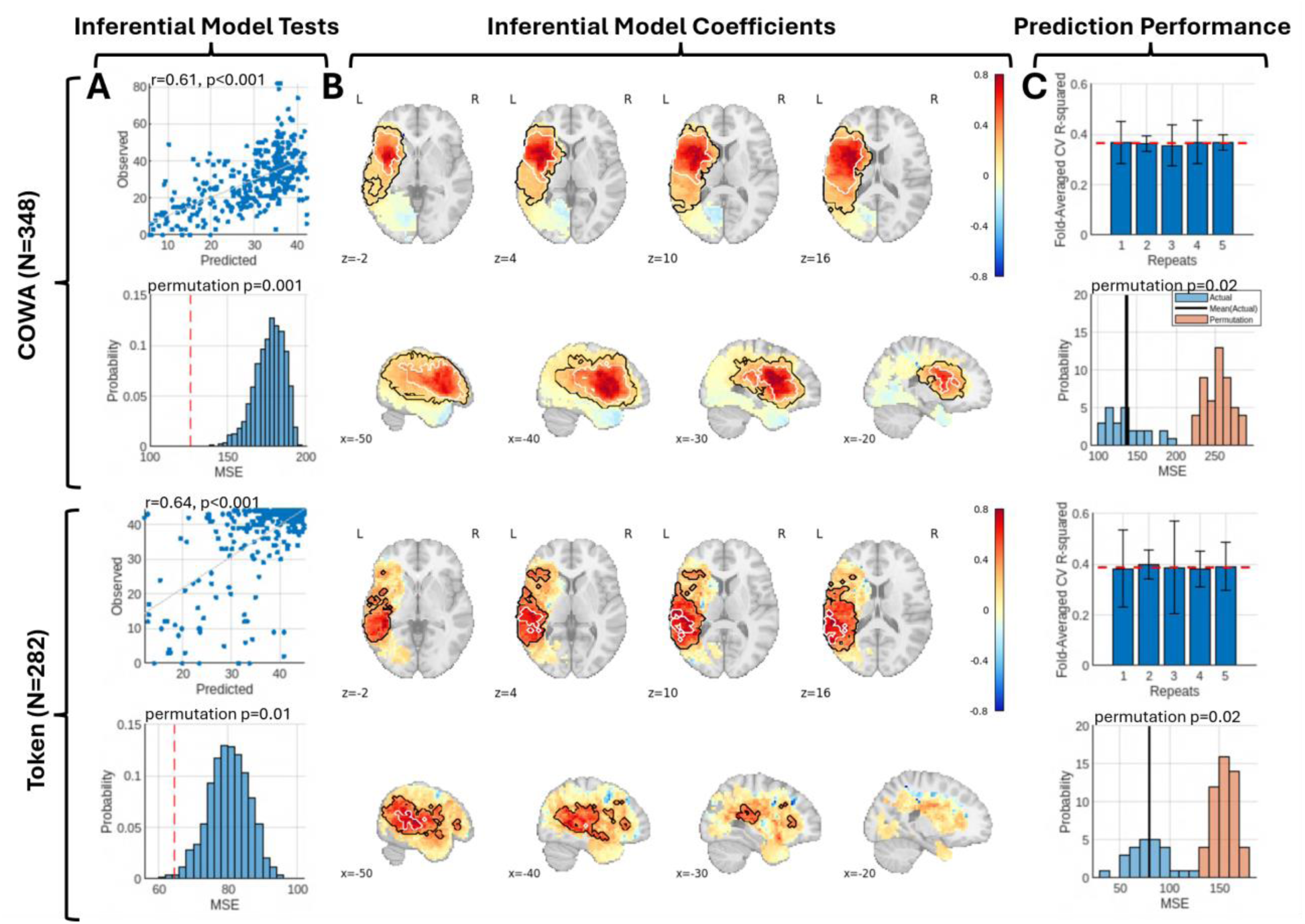
Multivariate lesion-behavior modeling of expressive and receptive language impairments **A.** Model-level tests evaluate whether there is a significant relationship between lesion location and behavior for the inferential models. The scatterplots show the full sample cross-validation correlations between average (across folds and repeats) out-of-fold predictions and observed scores for each outcome. The histograms show the actual observed MSE within the dataset (vertical dashed redline) relative to the permutation null distribution of MSE values. **B.** Coefficient-level results evaluate which regional brain-behavior relationships reach statistical significance within the inferential models. The unthresholded model coefficient maps are shown for each model. Voxels surviving at an FDR*p*<0.05 threshold are outlined in black, and voxels surviving an FWE*p*<0.05 threshold are outlined in white. **C.** Results of the predictive analyses estimate how much variance can be explained in held out data. The bar graphs show the fold-averaged cross-validation R^2^ values (and standard deviations) for each repetition of the cross-validation analyses. The horizontal dashed red lines indicate the average across all folds and repetitions. The histograms show the distribution of actual MSE scores across folds and repeats (blue histograms) along with the average MSE across all folds and repeats (black vertical line) and show the permutation null distribution of average (across folds and repeats) MSEs from the permutation tests on the cross-validation results (orange histograms). Note – model coefficients shown in (B) and (E) have been rescaled proportional to the maximum and minimum values in the maps.

#### Analysis 3: Multivariate Lesion-Behavior Classification

Next, to illustrate the application of the toolkit to categorical outcome data, we used PLS classification models to model the dichotomized outcome labels as a function of lesion location using the “pls_da” option in the toolkit. This analysis mirrored Analysis 2 but aimed to predict the binarized class labels corresponding to “impaired” (label=1) and “not impaired” (label=-1) patient sub-groups. By default, misclassification costs are equal for both groups. However, when there are group imbalances, modifying the misclassification costs can improve identification of the minority group. Here, misclassification costs for the “impaired” groups were set to reflect the difference in group representations in the dataset. For the COWA models, 42.82% of patients were assigned to the “impaired” group, and the misclassification cost for the “impaired” group was set to 1.3 times that of the “not impaired” group. For the Token Test models, 29.79% of patients were assigned to the “impaired” group, and the misclassification cost for the “impaired” group was set to 2.3 times that of the “not impaired” group. Predictions from PLS classification models are continuously varying prediction scores that must be dichotomized into group labels, and the default dichotomization approach is to take the sign of the predictions. Here, we used the prediction scores and class labels to construct an ROC curve. Then, we used the misclassification cost matrix and the ROC curve to identify optimal score cut-offs for dichotomization. For predictive analyses, the optimal score cut-offs were identified using the training folds and then applied to the prediction scores from the test folds to avoid data leakage. Otherwise, model training and statistical testing mirrored the approach described for Analysis 2.

The results of these analyses are summarized in **Figure 5**. Model-level tests of the inferential models using both the full sample cross-validation Fisher’s Exact Test on the confusion matrices constructed from the mode out-of-fold predicted class labels (**Figure 5A** – confusion matrices) and permutation tests of the inferential models fit to the full patient samples (**Figure 5A** – histograms) yielded statistically significant results, supporting evaluation of the model coefficients. Inspection of the coefficient maps from the inferential models indicated that the lesion locations that predicted the “impaired” group label were largely consistent with those identified by the regression analyses of the continuous task performance scores from Analysis 2 (**Figure 5B** vs. **Figure 4B**). Relative to the regression results from Analysis 1, the voxels that survived FWE correction for the Token Test exhibited a small but significant anterior and dorsal shift from the posterior temporal lobe to the posterior sylvian fissure (**Figure 5B** vs. **Figure 4B**), although the voxels that survived FDR correction were largely consistent across the two analyses. Notably, these results were highly consistent with the results of mass-univariate logistic regression analyses (**Supplemental Figure 1**). For the predictive analyses, both models had average out-of-sample AUCs of ∼0.8, which were statistically significant as indicated by the permutation tests of the full cross-validation procedures (**Figure 5C**). Similar prediction performance was observed for other types of multivariate models, and as for regression models, the use of DTLVC generally improved prediction performance for most models evaluated relative to not using DTLVC (**Supplementary Figures 4-5, 8-9**).

**Figure 5.**
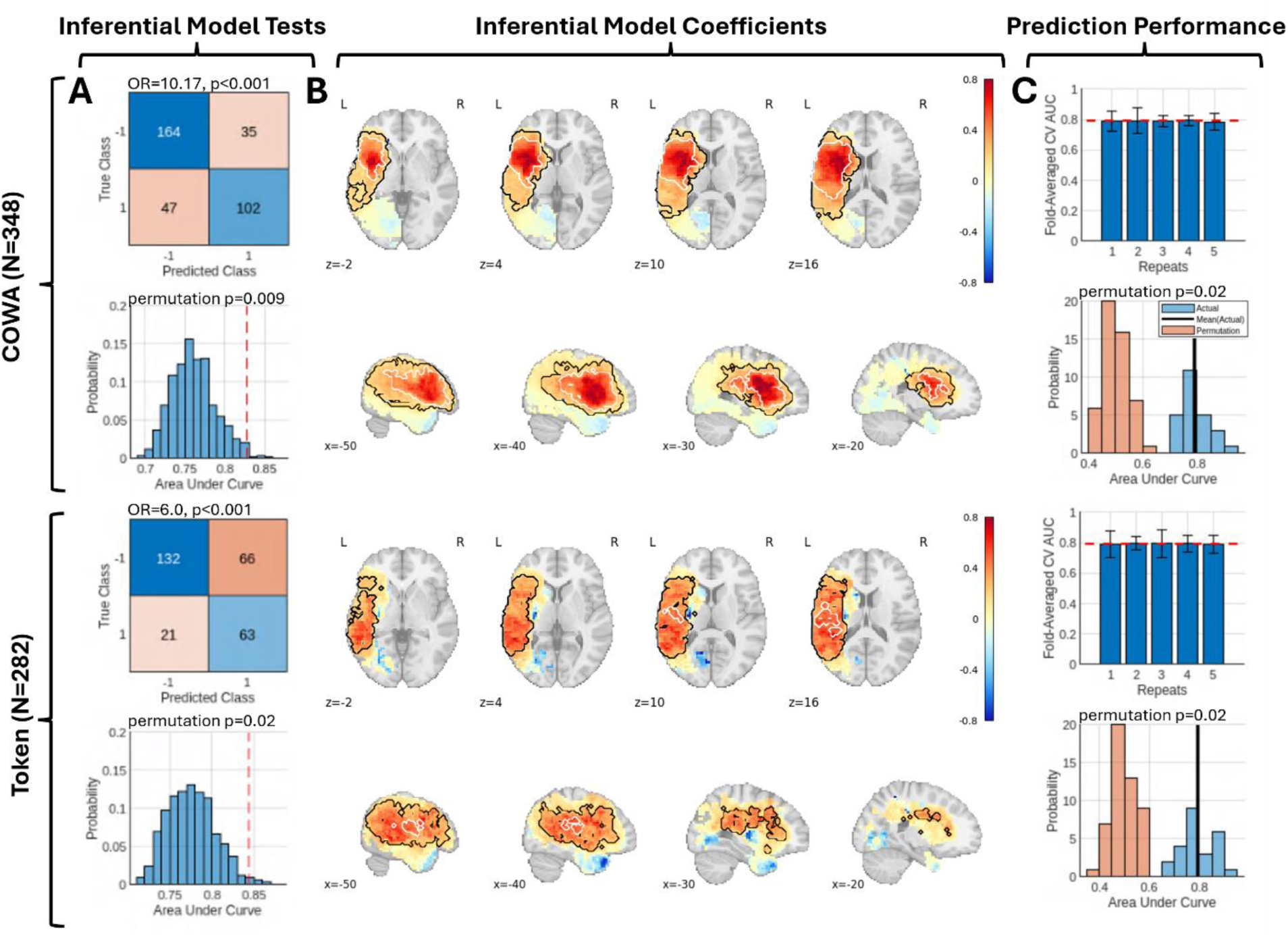
Multivariate lesion-behavior classification of expressive and receptive language impairments. **A.** Model-level tests for the inferential models. The confusion matrices show the full sample cross-validation classification results using the mode (across folds and repeats) out-of-fold predictions, along with the odds ratios (OR) and *p*-values from the Fisher’s Exact Tests. The histograms show the permutation null distributions of ROC AUCs for the inferential models fit to the full dataset, and the vertical dashed red lines indicate the observed AUCs of the inferential models. **B.** Coefficient-level results for the inferential models. The unthresholded model coefficient maps are shown each model. Voxels surviving at an FDR*p*<0.05 threshold are outlined in black, and voxels surviving an FWE*p*<0.05 threshold are outlined in white. **C.** Results of the predictive analyses. The bar graphs show the fold-averaged cross-validation R^2^ values (and standard deviations) for each repetition of the cross-validation analyses. The horizontal dashed red lines indicate the average across all folds and repetitions. The histograms show the distribution of actual AUC scores across folds and repeats (blue histograms) along with the average AUC across all folds and repeats (black vertical line), and show the permutation null distribution of average (across folds and repeats) AUCs from the permutation tests on the cross-validation results (orange histograms). Note – model coefficients shown in (B) and (E) have been rescaled proportional to the maximum and minimum values in the maps.

These results demonstrate the application of a classification-based approach to lesion-behavior modeling. Together with the results from Analysis 2, these results support the conclusion that the strongest lesion predictors of impairments on the COWA and Token test tend to localize to distinct lateral prefrontal and posterior temporal structures, respectively, but also indicate that voxels throughout the left MCA territory are relevant for discriminating between impaired vs. non-impaired patients. They also indicate that models trained on voxel-based lesion data can achieve relatively good performance at discriminating between impaired vs. unimpaired patients when applied to unseen patient samples, complementing the regression results reported in Analysis 2.

#### Analysis 4: Multivariate Disconnectome-Behavior Regressions

Next, to illustrate the application of a multivariate modeling approach to whole-brain disconnectomes, we performed multivariate regression analyses using the structural disconnection estimates as predictors. These analyses were performed identically to Analysis 2 (i.e., PLS regression with lesion predictors), but the model predictors corresponded to parcel-to-parcel disconnection matrices obtained by embedding each patient’s lesion into the normative tractography dataset; the DTLVC method for lesion volume correction was therefore not applied. Only edges (i.e., connections) with disconnection severity values of at least 25% in 10 or more patients were included in these analyses.

The results are summarized in **Figure 6**. Model-level tests of the inferential models for both the COWA and Token Test yielded highly significant full sample cross-validation correlations (**Figure 6A** – scatterplots) and permutation tests (**Figure 6A** -histograms), supporting further evaluation of the model coefficients. Inspection of the coefficient maps indicated that poor performance on the both the COWA and Token Test was associated widespread disconnection involving left frontal, temporal, and subcortical regions (**Figure 6B**), consistent with the results of the mass-univariate analyses (**Figure 3B**). It is worth noting that some of the connections with the largest weight magnitudes in the FDR-thresholded maps did not survive FWE correction (**Figure 6B**), highlighting that the strongest predictors in a model fit to the full sample may not be the most stable predictors in the model. This may be exacerbated by spatial biases in group-level lesion frequencies, as lesions affecting frontal regions were most common in these patient samples (**Figure 2**). The predictive analyses indicated that both models explained ∼35-40% of variance on average when applied to unseen patient samples, which was statistically significant as indicated by the permutation tests of the full cross-validation procedure (**Figure 6C**).

**Figure 6.**
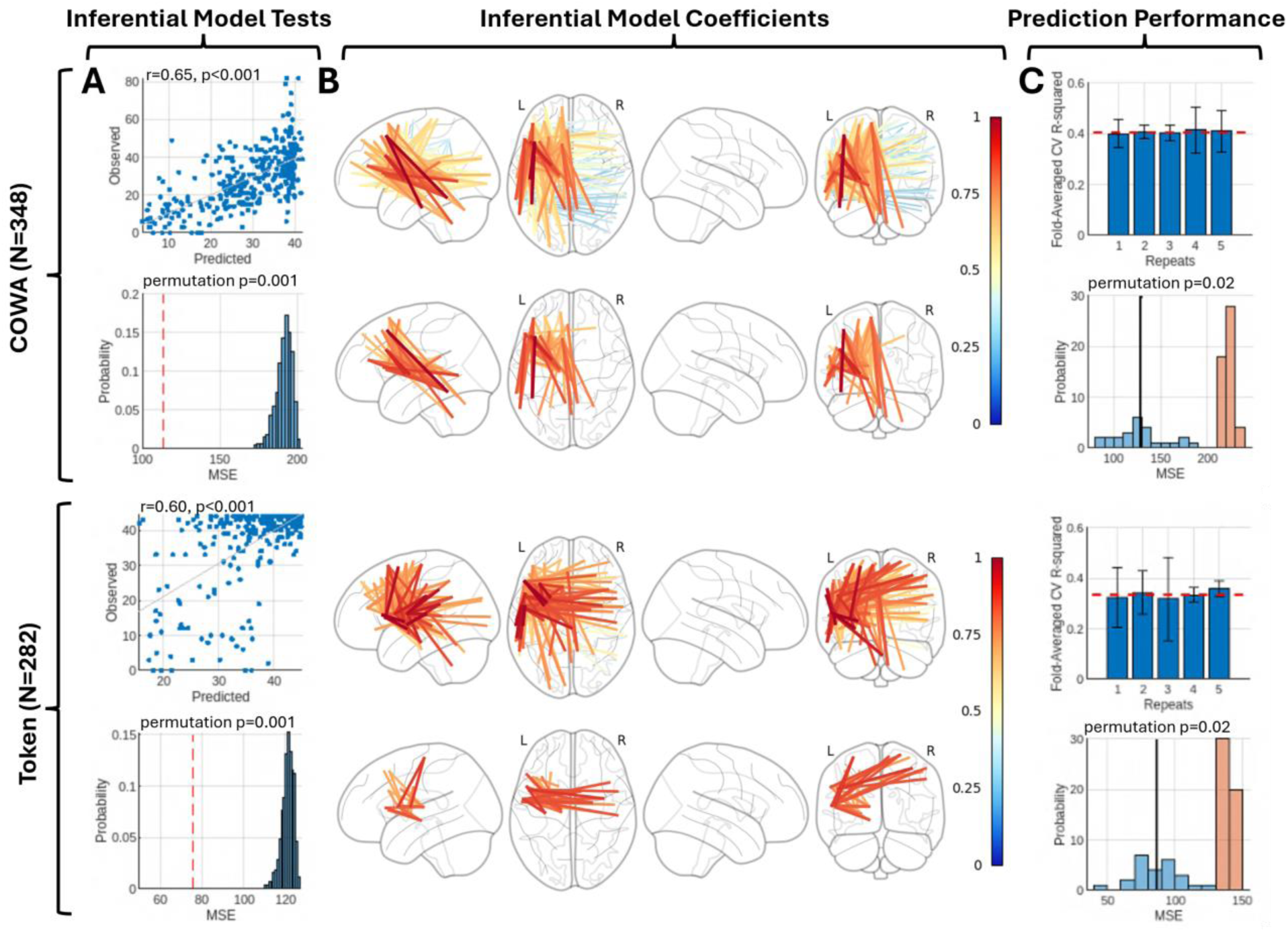
Multivariate disconnectome-behavior modeling of expressive and receptive language impairments. **A.** Model-level tests for the inferential models. The scatterplots show the full sample cross-validation correlations between average (across folds and repeats) out-of-fold predictions and observed scores for each outcome. The histograms show the permutation null distribution of MSE values for the inferential models fit to the full datasets, and the vertical dashed red lines indicate the observed MSEs of the inferential models fit to the full datasets. **B.** Coefficient-level results for the inferential models. The unthresholded model coefficient maps are shown each model. Voxels surviving at an FDR*p*<0.05 threshold are outlined in black, and voxels surviving an FWE*p*<0.05 threshold are outlined in white. **C.** Results of the predictive analyses. The bar graphs show the fold-averaged cross-validation R^2^ values (and standard deviations) for each repetition of the cross-validation analyses. The horizontal dashed red lines indicate the average across all folds and repetitions. The histograms show the distribution of actual MSE scores across folds and repeats (blue histograms) along with the average MSE across all folds and repeats (black vertical line), and show the permutation null distribution of average (across folds and repeats) MSEs from the permutation tests on the cross-validation results (orange histograms). Note – model coefficients shown in (B) and (E) have been rescaled proportional to the maximum and minimum values in the maps.

These results indicate that the strongest disconnection predictors of impairments on the COWA and Token Test map to partially distinct patterns of fronto-temporal and cortico-subcortical structural disconnections. They also suggest that models trained on parcel-to-parcel disconnection data perform similarly to models trained on voxel-based lesion data for predicting language outcomes. Speculatively, since structural disconnection data are likely most useful when they can capture common effects of non-overlapping lesions (Sperber et al., 2022), the observation that they perform similarly to voxel-based lesion data in these analyses may reflect the relatively dense lesion coverage in this sample (**Figure 2**) along with the relatively strict “minimum affection” threshold used to select voxels for inclusion in the models (i.e., only voxels with damage in at least 10 patients). Further work is needed to determine if/when derived measures such as structural disconnections will improve prediction performance, and this is an important avenue for future studies.

#### Analysis 5: Multivariate Functional Lesion Network-Behavior Regression

Next, to illustrate the application of a multivariate modeling approach to continuous whole-brain statistical maps, we performed multivariate regression analyses using the fLNM maps as predictors.

These analyses mirrored those in Analyses 4, except that the predictor data corresponded to functional lesion-network maps obtained from using each patient’s lesion as a seed in resting-state functional connectivity analyses in a normative sample. Only voxels with absolute fLNM *t*-statistics greater than 5 in at least 10 patients were included in the analyses.

The results of these analyses are shown in **Figure 7**. Model-level tests of the inferential models for both the COWA and Token Test yielded significant full-sample cross-validation correlations (**Figure 7A,D** – scatterplots) and significant permutation tests (**Figure 7A** – histograms), supporting further evaluation of the model coefficients. Inspection of the model coefficients indicated that impairments on both the COWA and Token Test were associated with lesions exhibiting functional connectivity to a left-lateralized fronto-temporal-parietal network (**Figure 7B**), consistent with the results of the mass-univariate analyses (**Figure 4C**). Notably, with FWE correction, the map for the Token Test primarily featured weights only in the bilateral posterior superior temporal cortex, suggesting that the local connectivity near the critical sites identified in Analysis 2 was most reliably predictive of impairment. In contrast, the FWE-corrected map for COWA included significant weights in both frontal and temporal cortices, suggesting potentially greater contributions of regions distal to the lesion sites identified in Analysis 2. The predictive analyses revealed that models trained on fLNM maps explained ∼25-30% of the variance on average when applied to unseen patient samples, which was statistically significant based on the permutation tests (**Figure 7C**). Notably, the models trained on the fLNM maps performed more poorly at predicting language impairments than the models trained on voxel-based lesion maps or parcel-to-parcel structural disconnection matrices (**Figure 4C**; **Figure 6C**), consistent with the results of previous studies comparing structural vs. functional lesion-derived networks for predicting outcomes in patients with brain lesions (Pini et al., 2021; Salvalaggio et al., 2020) Speculatively, this could reflect the mixing of signals from functionally distinct regions within the lesion mask seed regions, although it could also indicate that information about a lesion’s impact on brain structure is more predictive of language impairments than information about the functional connectivity patterns associated with the lesion. Further work is necessary to clarify if and/or when fLNM data can improve outcome prediction.

**Figure 7.**
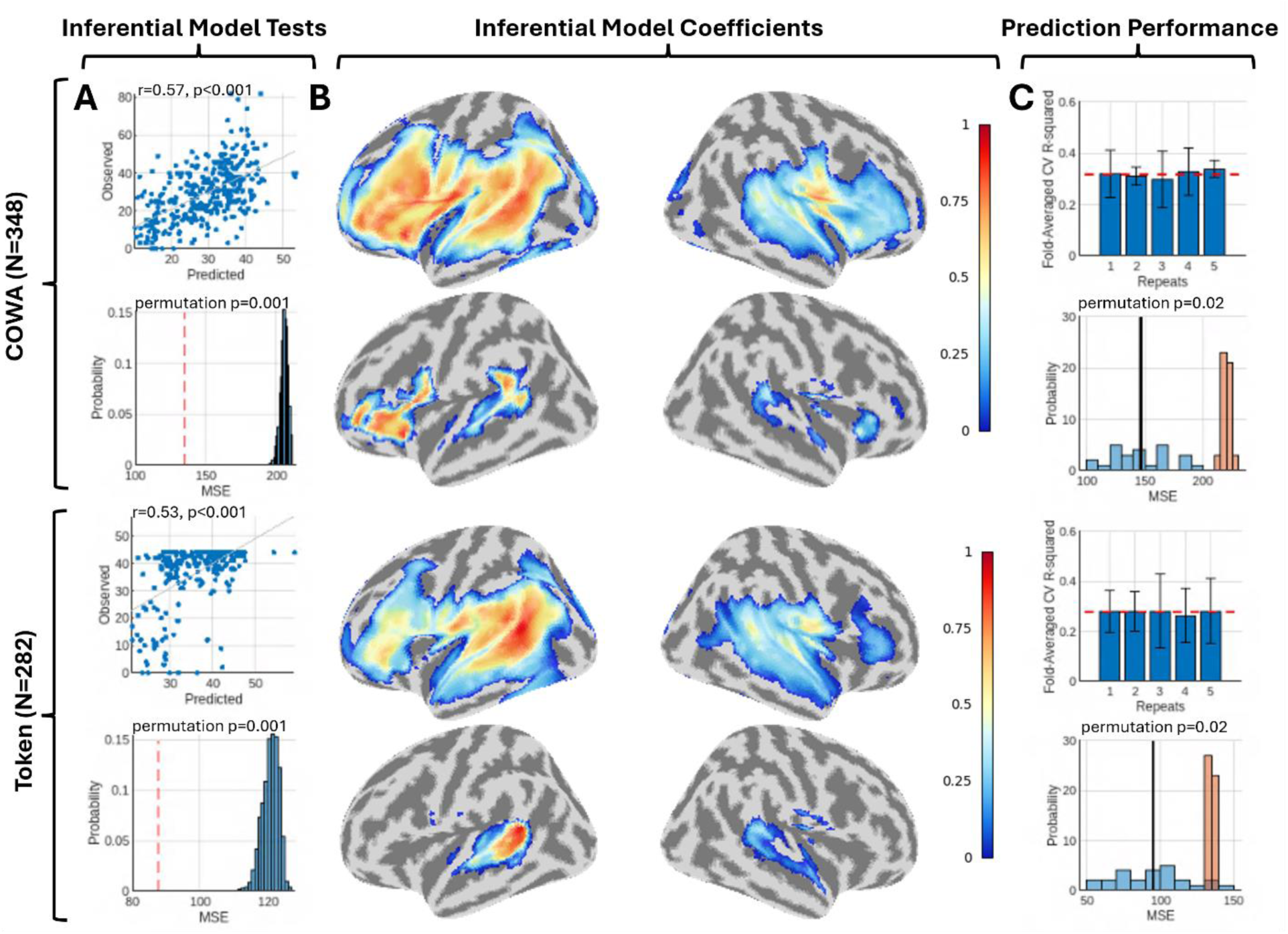
Multivariate functional network-behavior modeling of expressive and receptive language impairments. **A.** Model-level tests for the inferential models. The scatterplots show the full sample cross-validation correlations between average (across folds and repeats) out-of-fold predictions and observed scores for each outcome. The histograms show the permutation null distribution of MSE values for the inferential models fit to the full datasets, and the vertical dashed red lines indicate the observed MSEs of the inferential models fit to the full datasets. **B.** Coefficient-level results for the inferential models. The unthresholded model coefficient maps are shown each model. Voxels surviving at an FDR*p*<0.05 threshold are outlined in black, and voxels surviving an FWE*p*<0.05 threshold are outlined in white. **C.** Results of the predictive analyses. The bar graphs show the fold-averaged cross-validation R^2^ values (and standard deviations) for each repetition of the cross-validation analyses. The horizontal dashed red lines indicate the average across all folds and repetitions. The histograms show the distribution of actual MSE scores across folds and repeats (blue histograms) along with the average MSE across all folds and repeats (black vertical line) and show the permutation null distribution of average (across folds and repeats) MSEs from the permutation tests on the cross-validation results (orange histograms). Note – model coefficients shown in (B) and (E) have been rescaled proportional to the maximum and minimum values in the maps.

#### Analysis 6: Model Stacking

Recent work indicates that model stacking, either via averaging predictions across models trained on different predictor features (Olafson et al., 2023) or via training a “meta-model” to the predictions of different individual models (Pustina et al., 2017b) may improve predictive performance. We performed prediction averaging to combine the predictions of the lesion-behavior and disconnection-behavior regression models. This was performed by re-running the predictive cross-validation analyses and, within each outer loop, averaging the predictions of the out-of-fold observations across the two models. Results are shown in **Figure 8**. For COWA, there was no improvement in R^2^ with stacking of the models (**Figure 8A**). For Token Test, there was a very small increase in performance relative to the best performing base model, but this increase was not statistically significant (signed rank test: *p*=0.93). More complex model stacking approaches, such as fitting a random forest to the predictions of the base models, did not provide an advantage over model averaging (not shown). These results suggest that the different predictor modalities used here largely capture redundant behaviorally relevant information in these patient samples. Ultimately, further work is needed to determine how to best leverage model stacking to improve prediction performance.

**Figure 8.**
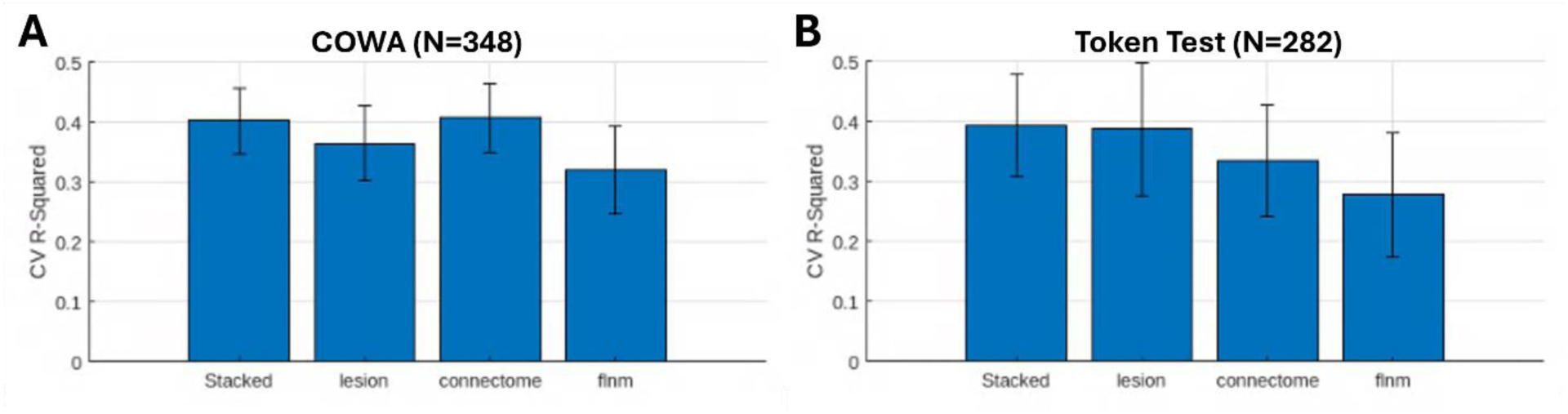
Model stacking results. **A.** The bar plots show the means and standard deviations of out-of-fold R-squared (y-axis) distributions for the stacked models, lesion models, structural disconnection models, and fLNM models (x-axis). The results for COWA are shown on the left, and the results for the Token Test are shown on the right.

## 4. Discussion

### Toolkit Summary

The methodology of lesion research has continued to evolve over the last 150 years, and progress has been especially rapid in recent years regarding the various statistical approaches available for relating brain lesions and behavioral outcomes (Moore et al., 2024). Multivariate lesion-symptom mapping has overcome many limitations that were present in earlier voxel-level analyses, and the integration of lesion mapping with human connectome data has facilitated our understanding of how lesions disrupt brain networks (Boes et al., 2015; Griffis et al., 2021; Sperber et al., 2022; Turken and Dronkers, 2011).

Here, we described a new software tool for inferential and predictive modeling of neuroimaging datasets, with a particular focus on modeling relationships between behavioral outcomes and neuroimaging-derived lesion measures. Importantly, this toolkit features a diverse set of tools for both inferential and predictive modeling, and features both a GUI and a scripting interface. It includes functionality for both mass-univariate and multivariate modeling strategies, includes implementations of both classification and regression models, and can flexibly accommodate different predictor modalities including NIFTI-formatted imaging data, adjacency matrices, and arbitrary predictor arrays. We anticipate that this will support comprehensive analysis approaches and facilitate systematic comparisons of different modeling strategies, predictor modalities, and parameter choices (Moore et al., 2024). Further, it automatically saves out key analysis parameters and generates “boilerplate” text descriptions of key methodological details and modeling results (Supplementary Material) to help ensure that methods and results are preserved even if the initial script that generated them is modified or lost. Overall, we hope that this tool will help to enhance the rigor and reproducibility of lesion research. We also hope that it will help to lay the groundwork for the eventual translation of research from this domain into future clinical applications that require dedicated predictive models, such as prognostication after a focal brain injury.

#### Lesion Correlates of Impairments on the COWA and Token Test

We also performed a series of example analyses in relatively large samples of patients with left hemisphere brain lesions. We used real imaging and behavioral data for tests of expressive and receptive language (i.e., COWA and Token Test) to demonstrate the application of the tool to real-world analysis applications. First, we performed mass-univariate analyses and fit inferential multivariate models using the full patient samples to identify lesion locations, structural disconnections, and lesion-derived functional networks associated with impaired language task performance.

Impaired performance on the COWA and other category fluency tests has previously been associated with damage to frontal, temporal, and parietal regions (Baldo et al., 2006; Biesbroek et al., 2015; Thye et al., 2020), although the results have not always been consistent across studies. Notably, one study reported that lesion sites associated with both semantic and phonemic fluency were largely confined to the left lateral prefrontal cortex (Biesbroek et al., 2015), which is largely consistent with our results. However, other studies have reported associations between fluency test performance and lesions to temporal areas as well (Baldo et al., 2006; Thye et al., 2020), but we did not observe significant associations between damage to temporal areas and impairments on the COWA. Rather, both our mass-univariate and multivariate analyses implicated a broad swath of left lateral prefrontal cortex including the inferior frontal gyrus, the insula, deep white matter, and subcortical structures (**Figure 3**, **Figure 4**, **Figure 5**). Even so, the analyses using structural disconnection matrices implicated fronto-temporal disconnections (**Figure 6**), and the analyses using fLNMs also implicated a lateral fronto-temporal functional network (**Figure 7**), indicating that fronto-temporal interactions are likely involved in performance on the COWA.

Impaired performance on the Token Test has previously been associated with lesions affecting the left posterior superior/middle temporal cortex and underlying white matter along with the inferior frontal gyrus pars triangularis (Adezati et al., 2022; Goldenberg et al., 2007), as well as to disrupted left temporal/parietal metabolism (Karbe et al., 1989). Our results are largely consistent with these observations, as they primarily implicate left temporal and parietal cortex, the underlying white matter, and to a lesser extent areas near the pars triangularis and ventral motor cortex (**Figure 3**, **Figure 4**, **Figure 5**). Our results also indicate that impairments on the Token Test are associated with structural disconnections affecting prefrontal, temporal, and subcortical regions (**Figure 6**), and with damage to a functional network featuring inferior frontal and posterior temporal/parietal cortices (**Figure 7**).

#### Predictive Modeling of the COWA and Token Test

We also performed predictive modeling analyses using in-sample cross-validation to evaluate the predictive power of models trained using lesion location, structural disconnection, and lesion-derived functional network features as predictors. Importantly, lesion-based, disconnection-based, and fLNM-based models were all able to achieve relatively good prediction performance on the COWA, explaining ∼37%, ∼40% and ∼32% of the variance in COWA performance on average when applied to unseen patient samples, respectively. However, combining these models via model stacking did not improve performance (**Figure 8**), suggesting that the lesion location and fLNM models did not capture unique information beyond that captured by the structural disconnection model. Lesion-based, disconnection-based, and fLNM-based models were also all able to achieve relatively good performance for the Token Test, explaining ∼38%, ∼34%, and ∼28% of the variance in Token Test performance when applied to unseen patient samples, respectively. In general, DTLVC tended to improve the predictive performance of multivariate models (**Supplemental Figures 3-9**), but more work is needed to systematically evaluate the impact of predictor transformations such as DTLVC on model performance. Combining the lesion, structural disconnection, and fLNM models via model stacking resulted in a small improvement in performance for the Token Test (**Figure 8B**), with the stacked model explaining ∼1% more variance on average when applied to unseen patient samples. However, stacking did not improve prediction performance for COWA (**Figure 8A**). Overall, there was not a substantial advantage to performing model stacking in these analyses.

In general, the individual models showed quite good performance that was comparable to or better than what has been reported for similar lesion-derived predictive models in the literature (Corbetta et al., 2015; Olafson et al., 2023; Salvalaggio et al., 2020; Talozzi et al., 2023; Yourganov et al., 2016), even though some of these models used strategies that could optimistically bias results (Poldrack et al., 2020; Scheinost et al., 2019) such as full-sample feature selection, leave-one-out CV, and correlation as the measure of prediction performance (Corbetta et al., 2015; Yourganov et al., 2016). While we did not observe a substantial benefit of model stacking in these analyses, it is possible that stacking a larger set of models trained on a more diverse set of features would further improve model performance (Pustina et al., 2017b). However, it is worth noting that other work has reported similarly modest increases in performance when stacking models to predict post-stroke motor outcomes (Olafson et al., 2023). Ultimately, more work is needed to determine how to best leverage model stacking to improve predictive performance. By making our toolkit available to other researchers, we hope to facilitate advances towards identifying optimal strategies for predicting outcomes after brain injury.

#### Recommendations and Future Directions

While providing detailed recommendations about the specific modeling approaches, parameter choices, and sample sizes that should be used for specific analyses is beyond the scope of the current study, we nonetheless urge potential users to carefully consider these factors when designing their study and specifying their modeling approaches.

In general, sample sizes of at least 100 have been recommended for inferential applications of multivariate models in lesion symptom-mapping studies, and mass-univariate methods may be better-suited for inferential analyses in smaller patient samples (Ivanova et al., 2021b; Sperber et al., 2018); the permutation-based continuous family-wise error control procedure, which the toolkit uses by default for mass-univariate approaches, has in particular been shown to outperform other multiple comparisons corrections methods in samples of 30-60 patients and is therefore recommended for inferential analyses of small patient samples (Mirman et al., 2018). For predictive modeling analyses, the minimum sample sizes recommended by previous studies range from one hundred to several hundred patients (e.g.,

Scheinost et al., 2019; Poldrack et al., 2020). In general, predictive model performance is expected to improve as sample size increases (e.g., see Figure 4 of Scheinost et al., 2019), but it is important to note that this is only likely to be true if the patient characteristics are appropriate given the goals of the analysis and assumptions of the approach (Scheinost et al., 2019; Moore et al., 2024).

Indeed, whether a given sample size is sufficient to allow for the detection of effects and/or the development of robust models will depend on the spatial properties of the lesion data and the distribution of deficits in the sample under study, since this will determine the effective sample size and power at each voxel or predictor variable (see Figure 3 in Moore et al., 2024). For example, a sample of 300 patients may still be poorly suited for lesion-symptom analyses if only 3 or 4 patients have overlapping lesions at any given voxel, or if there is little-to-no variability in behavioral scores between patients with vs. without damage to sufficiently powered voxels. Minimum lesion affection thresholds should also be selected carefully (Sperber et al., 2016), and users should ensure that they are appropriate for the modeling strategy employed; robust effect estimates cannot be obtained for voxels that are only damaged in one or two patients. In general, care should be taken to ensure that the sample used to train and evaluate a model is reasonably representative of the population of interest (i.e., the population to which the model is intended to generalize). For example, performance estimates for a model trained to predict a left-lateralized function (e.g., language deficits) in a sample of patients with equal numbers of left unilateral and right unilateral lesions may be optimistically biased if much of the total sample variance in task performance could be attributed to lesion laterality --if right unilateral lesion patients consistently perform at ceiling while left hemisphere patients do not, then the model might assign a perfect score to every patient with lesions to the right of the midline, resulting in very small prediction errors for half of the patient sample (i.e., for patients with right unilateral lesions). The resulting performance estimates could appear much better than performance estimates obtained from a model trained only using data from the subset of patients with left unilateral lesions (i.e., the population of interest), even if the performance for the left unilateral lesion patients were identical for both models. In such a case, the performance estimates obtained at the level of the full sample may not be reflective of the performance of the model for the true population of interest (i.e., patients with left hemisphere lesions). It is therefore important to ensure that the patient sample is reflective of the population of interest given the hypotheses and/or intended model applications. Additionally, it is worth noting that increasing the sample size can increase the likelihood that very weak effects will achieve statistical significance in inferential analyses, highlighting the importance of incorporating information about effect size when interpreting statistically significant effects (Lorca-Puls et al., 2018). For general guidelines on lesion analysis, we refer readers to the recent paper by Moore et al., (2024). For guidelines on predictive modeling in the context of neuroimaging, we refer readers to the papers by Poldrack et al., (2020) and Scheinost et al., (2019).

As shown in **Table 1**, the IBB Toolkit provides diverse functionality compared to existing toolkits. Systematic comparisons of the different modeling techniques, analysis options, and parameter choices available in the toolkit to identify optimal strategies for lesion-behavior modeling are an important avenue for future work. This could be accomplished using simulation studies and applications to real world data. Importantly, the toolkit provides a platform for conducting rigorous methodological comparisons, and we hope that this will facilitate the identification of optimal strategies and development of evidence-based guidelines for the development, refinement, and testing of lesion-behavior models.

## 5. Conclusions

Here, we presented a new open-source software tool for inferential and predictive modeling of neuroimaging and lesion data. We have made this tool freely available to the research community with the goal of facilitating advanced modeling analyses of lesion and other neuroimaging datasets. To this end, we have also included a set of tutorial MATLAB Live notebooks that walk users through a range of different analyses using the toolkit and explain each step and the associated inputs, options, and result outputs. In addition, we provide a dedicated User Manual that provides step-by-step instructions for performing modeling analyses using the GUI. The toolkit, including the tutorial notebooks, can be downloaded under a GNU GPL v3.0 license at (https://github.com/jcgriffis/ibb_toolkit).

## Acknowledgements

All procedures performed in the study were in accordance with the University of Iowa Institutional Review Board. This study was supported by the National Institute of Neurological Disease and Stroke (1 R01 NS114405), the Roy J. Carver Trust (J.C.G and A.D.B), the University of Iowa Medical Scientist Training Program (S.F.A, funded by 5T32GM139776-04) and the University of Iowa Department of Pediatrics. This work was conducted on an MRI instrument funded by 1S10OD025025-01.

## 7. Author Contributions

J.C.G conceptualized the toolkit, developed and tested the toolkit software, performed data processing, designed and implemented the analyses, and drafted and edited the manuscript. J.B. processed the lesion and functional lesion-network data and provided computing support. S.F.A. supported toolkit development and testing and contributed to editing the manuscript. C.S. participated in data collection and provided support with identifying and organizing behavioral, clinical, and demographic data. D.T. provided data and other resources. A.D.B. provided supervision, funding, and other resources, and contributed to drafting and editing the manuscript.

## 8. Data Availability

The datasets generated and/or analyzed during the current study are available upon reasonable request to the corresponding author. The toolkit used to run the analyses reported in the paper is available at https://github.com/jcgriffis/ibb_toolkit.

## References

Abdi, H. (2010). Partial least squares regression and projection on latent structure regression (PLS Regression). Wiley Interdiscip. Rev. Comput. Stat. 2, 97–106.

Adezati, E., Thye, M., Edmondson-Stait, A.J., Szaflarski, J.P., and Mirman, D. (2022). Lesion correlates of auditory sentence comprehension deficits in post-stroke aphasia. Neuroimage: Reports 2, 100076.

Aickin, M., and Gensler, H. (1996). Adjusting for multiple testing when reporting research results: The Bonferroni vs Holm methods. Am. J. Public Health 86, 726–728.

Baldo, J. V, Schwartz, S., Wilkins, D., and Dronkers, N.F. (2006). Role of frontal versus temporal cortex in verbal fluency as revealed by voxel-based lesion symptom mapping. J. Int. Neuropsychol. Soc. 12, 896– 900.

Benjamini, Y., and Hochberg, Y. (1995). Controlling the False Discovery Rate: A Practical and Powerful Approach to Multiple Testing. J. R. Stat. Soc. Ser. B 57, 289–300.

Biesbroek, J.M., van Zandvoort, M.J.E., Kappelle, L.J., Velthuis, B.K., Biessels, G.J., and Postma, A. (2015). Shared and distinct anatomical correlates of semantic and phonemic fluency revealed by lesion-symptom mapping in patients with ischemic stroke. Brain Struct. Funct.

Boes, A., Prasad, S., Pascual-Leone, A., H, L., Q, L., Caviness, V., and Fox, M.D. (2015). Network localization of neurological symptoms from focal brain lesions. Brain, Press. 3061–3075.

Bowren, M., Bruss, J., Manzel, K., Edwards, D., Liu, C., Corbetta, M., Tranel, D., and Boes, A.D. (2022). Post-stroke outcomes predicted from multivariate lesion-behaviour and lesion network mapping. Brain 145, 1338–1353.

Bzdok, D., and Yeo, B.T.T. (2017). Inference in the age of big data: Future perspectives on neuroscience. Neuroimage 155, 549–564.

Corbetta, M., Ramsey, L., Callejas, A., Baldassarre, A., Hacker, C.D., Siegel, J.S., Astafiev, S.V., Rengachary, J., Zinn, K., Lang, C.E., et al. (2015). Common Behavioral Clusters and Subcortical Anatomy in Stroke. Neuron 85, 927–941.

Damasio, H., and Damasio, A.R. (1989). Lesion Analysis in Neuropsychology (Oxford University Press).

DeMarco, A.T., Turkeltaub, P.E., and DeMarco, Andrew; Turkeltaub, P. (2018). A multivariate lesion symptom mapping toolbox and examination of lesion-volume biases and correction methods in lesion-symptom mapping. Hum. Brain Mapp. 21, 2461–2467.

Diedrichsen, J. (2006). A spatially unbiased atlas template of the human cerebellum. Neuroimage 33, 127–138.

Friederici, A.D., and Gierhan, S.M.E. (2013). The language network. Curr. Opin. Neurobiol. 23, 250–254.

Goldenberg, G., Hermsdörfer, J., Glindemann, R., Rorden, C., and Karnath, H.-O. (2007). Pantomime of Tool Use Depends on Integrity of Left Inferior Frontal Cortex. Cereb Cortex.

Griffis, J.C., Nenert, R., Allendorfer, J.B., and Szaflarski, J.P. (2017). Damage to white matter bottlenecks contributes to language impairments after left hemispheric stroke. NeuroImage Clin. 14, 552–565.

Griffis, J.C., Metcalf, N. V, Corbetta, M., and Shulman, G.L. (2019). Structural disconnections explain brain network dysfunction after stroke. Cell Rep. 28, 2527–2540.

Griffis, J.C., Metcalf, N. V, Corbetta, M., and Shulman, G.L. (2021). Lesion Quantification Toolkit: A MATLAB software tool for estimating grey matter damage and white matter disconnections in patients with focal brain lesions. NeuroImage Clin. 30.

Ivanova, M. V., Herron, T.J., Dronkers, N.F., and Baldo, J. V. (2021a). An empirical comparison of univariate versus multivariate methods for the analysis of brain–behavior mapping. Hum. Brain Mapp. 42, 1070–1101.

Ivanova, M. V, Herron, T.J., Dronkers, N.F., and Baldo, J. V (2021b). An empirical comparison of univariate versus multivariate methods for the analysis of brain–behavior mapping. Hum. Brain Mapp. 42, 1070–1101.

Jiang, Y., and Gong, G. (2024). Common and distinct patterns underlying different linguistic tasks: multivariate disconnectome symptom mapping in poststroke patients. Cereb. Cortex 34, 1–11.

Joutsa, J., Corp, D.T., and Fox, M.D. (2022a). Lesion network mapping for symptom localization: Recent developments and future directions. Curr. Opin. Neurol. 35, 453–459.

Joutsa, J., Moussawi, K., Siddiqi, S.H., Abdolahi, A., Drew, W., Cohen, A.L., Ross, T.J., Deshpande, H.U., Wang, H.Z., Bruss, J., et al. (2022b). Brain lesions disrupting addiction map to a common human brain circuit. Nat. Med. 28, 1249–1255.

Karbe, H., Herholz, K., Szelies, B., Pawlik, G., Wienhard, K., and Heiss, W.D. (1989). Regional metabolic correlates of token test results in cortical and subcortical left hemispheric infarction. Neurology 39, 1083–1088.

Kohoutová, L., Heo, J., Cha, S., Lee, S., Moon, T., Wager, T.D., and Woo, C.W. (2020). Toward a unified framework for interpreting machine-learning models in neuroimaging. Nat. Protoc. 15, 1399–1435.

Lorca-Puls, D.L., Gajardo-Vidal, A., White, J., Seghier, M.L., Leff, A.P., Green, D.W., Crinion, J.T., Ludersdorfer, P., Hope, T.M.H., Bowman, H., et al. (2018). The impact of sample size on the reproducibility of voxel-based lesion-deficit mappings. Neuropsychologia 115, 101–111.

Mirman, D., Chen, Q., Zhang, Y., Wang, Z., Faseyitan, O.K., Coslett, H.B., and Schwartz, M.F. (2015). Neural organization of spoken language revealed by lesion–symptom mapping. Nat. Commun. 6, 6762.

Mirman, D., Landrigan, J.F., Kokolis, S., Verillo, S., Ferrara, C., and Pustina, D. (2017). Corrections for multiple comparisons in voxel-based lesion-symptom mapping. Neuropsychologia 115, 112–123.

Moore, M.J., Demeyere, N., Rorden, C., and Mattingley, J.B. (2024). Lesion mapping in neuropsychological research: A practical and conceptual guide. Cortex 170, 38–52.

Olafson, E.R., Sperber, C., Jamison, K.W., Bowren, M.D., Boes, A.D., Andrushko, J.W., Borich, M.R., Boyd, L.A., Cassidy, J.M., Conforto, A.B., et al. (2023). Data-driven biomarkers outperform theory-based biomarkers in predicting stroke motor outcomes. BioRxiv 16, 2023.06.19.545638.

Pini, L., Salvalaggio, A., De Filippo De Grazia, M., Zorzi, M., de Schotten, M.T., and Corbetta, M. (2021). A novel stroke lesion network mapping approach: improved accuracy yet still low deficit prediction. Brain Commun. 3.

Poldrack, R.A., Huckins, G., and Varoquaux, G. (2020). Establishment of Best Practices for Evidence for Prediction: A Review. JAMA Psychiatry 77, 534–540.

Pustina, D., Avants, B., Faseyitan, O., Medaglia, J., and Branch Coslett, H. (2017a). Improved accuracy of lesion to symptom mapping with multivariate sparse canonical correlations. Neuropsychologia 8000, 1– 13.

Pustina, D., Coslett, H.B., Ungar, L., Faseyitan, O.K., Medaglia, J.D., Avants, B., and Schwartz, M.F. (2017b). Enhanced estimations of post-stroke aphasia severity using stacked multimodal predictions. Hum. Brain Mapp. 00.

Rey, G.J., Feldman, E., Rivas-Vazquez, R., Levin, B.E., and Benton, A. (1999). Neuropsychological test development and normative data on Hispanics. Arch. Clin. Neuropsychol. 14, 593–601.

Rorden, C., and Karnath, H.-O. (2004). Using human brain lesions to infer function: a relic from a past era in the fMRI age? Nat. Rev. Neurosci. 5, 813–819.

Salvalaggio, A., Grazia, M.D.F. De, Schotten, M.T. De, Corbetta, M., and Zorzi, M. (2020). Post-stroke deficit prediction from lesion and indirect structural and functional disconnection. Brain.

Schaefer, A., Kong, R., Gordon, E.M., Laumann, T.O., Zuo, X.-N., Holmes, A.J., Eickhoff, S.B., and Yeo, B.T.T. (2018). Local-Global Parcellation of the Human Cerebral Cortex from Intrinsic Functional Connectivity MRI. Cereb. Cortex 28, 3095–3114.

Scheinost, D., Noble, S., Horien, C., Greene, A.S., Lake, E.M., Salehi, M., Gao, S., Shen, X., O’Connor, D., Barron, D.S., et al. (2019). Ten simple rules for predictive modeling of individual differences in neuroimaging. Neuroimage 193, 35–45.

Sperber, C., and Karnath, H.-O. (2016). Impact of correction factors in human brain lesion-behavior inference. Hum. Brain Mapp. 1701, 1692–1701.

Sperber, C., Wiesen, D., and Karnath, H.-O. (2018). An empirical evaluation of multivariate lesion behaviour mapping using support vector regression. Hum. Brain Mapp. hbm.24476.

Sperber, C., Griffis, J., and Kasties, V. (2022). Indirect structural disconnection-symptom mapping. Brain Struct. Funct. 227, 3129–3144.

Talozzi, L., Forkel, S.J., Pacella, V., Nozais, V., Allart, E., Piscicelli, C., Pérennou, D., Tranel, D., Boes, A., Corbetta, M., et al. (2023). Latent disconnectome prediction of long-term cognitive-behavioural symptoms in stroke. Brain 146, 1963–1978.

Thye, M., Szaflarski, J.P., and Mirman, D. (2020). Shared lesion correlates of semantic and letter fluency in post-stroke aphasia. J. Neuropsychol. 1–8.

Turken, A.U., and Dronkers, N.F. (2011). The neural architecture of the language comprehension network: converging evidence from lesion and connectivity analyses. Front. Syst. Neurosci. 5, 1.

Varoquaux, G., Raamana, P.R., Engemann, D.A., Hoyos-Idrobo, A., Schwartz, Y., and Thirion, B. (2017). Assessing and tuning brain decoders: Cross-validation, caveats, and guidelines. Neuroimage 145, 166– 179.

Woo, C.-W., Krishnan, A., and Wager, T.D. (2014). Cluster-extent based thresholding in fMRI analyses: pitfalls and recommendations. Neuroimage 91, 412–419.

Yeh, F.C. (2022). Population-based tract-to-region connectome of the human brain and its hierarchical topology. Nat. Commun. 13.

Yourganov, G., Fridriksson, J., Rorden, C., Gleichgerrcht, E., and Bonilha, L. (2016). Multivariate Connectome-Based Symptom Mapping in Post-Stroke Patients: Networks Supporting Language and Speech. J. Neurosci. 36, 6668–6679.

Yourganov, G., Fridriksson, J., and Rorden, C. (2018). Estimating the statistical significance of spatial maps for multivariate lesion-symptom analysis. Cortex 108, 276–278.

Zhang, Y., Kimberg, D.Y., Coslett, H.B., Schwartz, M.F., Wang, Z., Yongsheng, Z., Kimberg, D.Y., Coslett, H.B., Schwartz, M.F., and Wang, Z. (2014). Multivariate lesion-symptom mapping using support vector regression. Hum. Brain Mapp. 35, 997.

