## Supplemental Material for "Iowa Brain-Behavior Modeling Toolkit: An Open-Source MATLAB Tool for Inferential and Predictive Modeling of Imaging-Behavior and Lesion-Deficit Relationships"

**Supplemental Material 1. Automatically Generated Methods and Results Text for COWA**

**Mass-Univariate**

*Methods*

Data were analyzed using mass-univariate Pearson correlation analyses. Only predictors with at least 10 non-zero observations with absolute values greater than 1 were included in the analysis. After excluding observations with no data for any included predictors, N=328 observations were included in the analysis. An inferential model was fit to the full dataset. To determine the significance of the inferential model compared to an empirical null distribution of model fits, permutation testing was performed using 10000 permutation iterations. The continuous FWE method (Mirman et al., 2017) was used to determine a single family-wise error threshold across all predictors at predictor thresholds of [1,10,50,100,500,1000]. Coefficient-level permutation p-values were also generated. Bonferroni-Holm correction was used to obtain FWE-corrected coefficient-level permutation p-values. The Benjamini-Hochberg procedure was used to obtain FDR-corrected coefficient-level permutation p-values.

*Results*

N/A (no model results, only map outputs)

**PLS Regression Model**

*Methods*

Data were modeled using partial least squares regression (PLSR). Only predictors with at least 10 non-zero observations with absolute values greater than 1 were included in the analysis. N=348 observations were included in the analysis. The direct total lesion volume control method (Zhang et al., 2014) was used to mitigate the effects of lesion volume. An inferential model was fit to the full dataset. Hyper-parameter optimization was performed to determine the k parameter (number of PLS components) using 5 repetitions of 5-fold cross-validation with repartitioning at each repetition. To determine the significance of the inferential model compared to an empirical null distribution of model fits, permutation testing was performed using 1000 permutation iterations. Bootstrap resampling with 1000 bootstrap iterations was used to estimate the statistical significance of individual coefficients (Kohoutova et al., 2020). The Bonferroni-Holm procedure was used to obtain FWE-corrected coefficient-level p-values. The Benjamini-Hochberg procedure was used to obtain FDR-corrected coefficient-level p-values. 95% bootstrap confidence intervals were estimated for the model fit, and FWE-corrected 95% bootstrap confidence intervals were estimated for individual coefficients using the normal approximation method. Nested cross-validation was used to optimize model hyper-parameters and obtain estimates of out-of-sample prediction performance. Cross-validation analyses were performed using 5 repetitions of 5-fold cross-validation with repartitioning at each repetition. Train and test partitions were stratified according to the outcome variable. To test the significance of the inferential model in terms of the cross-validation performance, the average (across cross-validation repetitions) Pearson correlation was computed between the out-of-fold predictions and observed outcomes for all observations (Pustina et al., 2017). Out-of-sample prediction performance was estimated using the cross-validation explained variance score, prediction R-squared, Pearson correlation of predicted and observed outcomes, and mean squared error. Permutation testing was performed using 50 iterations of the full cross-validation procedure to determine the statistical significance of the cross-validation out-of-sample prediction performance. The average (across folds and repeats) MSE from the cross-validation run was compared against the null distribution obtained from the permutation analyses to determine statistical significance of the full cross-validation results.

*Results*

The inferential model fit to the full dataset explained 41.5286% of the variance in the outcome (95% CI: [0.30782, 0.45014]). The permutation test indicated that the model fit better than expected given the permutation null distribution (p=0.000999). The cross-validation test indicated that predicted out-of-fold outcomes were significantly correlated with the observed outcomes across the full dataset (r=0.60635, p=2.5845e-36), supporting the interpretation of the inferential model. Across all folds and repeats, the average cross-validation R-squared was 0.36533 (SD=0.062136). Permutation testing on the full cross-validation procedure indicated that the average cross-validation loss (MSE) was lower than expected given the permutation null distribution (p=0.019608).

**PLS Classification Model**

*Methods*

Data were modeled using partial least squares discriminant analysis (PLS-DA). Only predictors with at least 10 non-zero observations with absolute values greater than 1 were included in the analysis. N=348 observations were included in the analysis. The direct total lesion volume control method (Zhang et al., 2014) was used to mitigate the effects of lesion volume. An inferential model was fit to the full dataset. Hyper-parameter optimization was performed to determine the k parameter (number of PLS components) using 5 repetitions of 5-fold cross-validation with repartitioning at each repetition. To determine the significance of the inferential model compared to an empirical null distribution of model fits, permutation testing was performed using 1000 permutation iterations. Bootstrap resampling with 1000 bootstrap iterations was used to estimate the statistical significance of individual coefficients (Kohoutova et al., 2020). The Bonferroni-Holm procedure was used to obtain FWE-corrected coefficient-level p-values. The Benjamini-Hochberg procedure was used to obtain FDR-corrected coefficient-level p-values. 95% bootstrap confidence intervals were estimated for the model fit, and FWE-corrected 95% bootstrap confidence intervals were estimated for individual coefficients using the normal approximation method. Nested cross-validation was used to optimize model hyper-parameters and obtain estimates of out-of-sample prediction performance. Cross-validation analyses were performed using 5 repetitions of 5-fold cross-validation with repartitioning at each repetition. Train and test partitions were stratified according to the outcome variable. To test the significance of the inferential model in terms of the cross-validation performance, Fishers exact test was performed on the confusion matrix computed using the mode (across cross-validation repetitions) out-of-sample predicted classes and true outcomes for all observations (analogous to Pustina et al., 2017). Out-of-sample prediction performance was estimated using the cross-validation classification accuracy for each class, the overall classification accuracy, and the area under the ROC curve. Permutation testing was performed using 50 iterations of the full cross-validation procedure to determine the statistical significance of the cross-validation out-of-sample prediction performance. The average (across folds and repeats) area under the ROC curve from the cross-validation run was compared against the null distribution obtained from the permutation analyses to determine statistical significance of the full cross-validation results. The misclassification cost assigned to the class labeled -1 was set to 1, and the misclassification cost assigned to the class labeled 1 was set to 1.3.

*Results*

The inferential model fit to the full dataset achieved 84.4221% classification accuracy for the group labeled -1 (95% CI: [0.76371, 0.96639]), 72.4832% accuracy for the group labeled 1 (95% CI: [0.55488, 0.75676]), and 79.3103% accuracy for the combined groups (95% CI: [0.73687, 0.81314]). The area under the ROC curve for the inferential model fit to the full dataset was 0.82802 (95% CI: [0.76609, 0.83166]). The permutation test indicated that the inferential model fit the full dataset better than expected given the permutation null distribution (p=0.010989). The cross-validation test indicated that predicted out-of-fold outcomes were significantly associated with the observed outcomes (OR=10.169, p=2.8069e-22), supporting the interpretation of the inferential model. Cross-validation was used to estimate out-of-sample prediction performance. Across all folds and repeats, the average cross-validation area under the ROC curve was 0.79104 (SD=0.052757). Permutation testing on the full cross-validation procedure indicated that the average cross-validation area under the ROC curve was greater than expected given the permutation null distribution (p=0.019608).

**Supplementary Material 2. Lesion-Behavior Analyses Using Different Modeling Strategies**

We performed lesion-behavior analyses using different modeling strategies implemented in the toolkit. For the analyses in **Supplemental Figure 1**, we performed mass-univariate Brunner-Munzel tests with lesion volume regression, mass-univariate unequal variance t-tests with lesion volume regression, and mass-univariate linear regressions and logistic regressions (using the dichotomized outcome variables) with lesion volume included as a covariate in the models. These analyses used the same general parameters reported for the mass-univariate correlation analyses in the main text, although the voxel threshold *v* was increased from *v*=100 to *v*=1000 for the mass-univariate t-test analysis of the Token Test due to no voxels surviving at the *v*=100 threshold. Maps in **Supplemental Figure 1** show test statistics (i.e., t-statistics, Brunner-Munzel statistics).

For the analyses in **Supplemental Figures 2-9**, we performed multivariate regression and classification analyses using different models implemented in the toolkit. These analyses used the same general analysis parameters as the analyses reported in the main text, with the exception that the outcome variable was standardized for the “rbfsvr” (i.e., SVR with radial basis function kernel) models (in our experience, this improves performance for this model). Maps in **Supplemental Figures 2-9** show model coefficient magnitudes rescaled as proportion of the maximum value in each map for all models except for “rbfsvr” and “rbfsvc”, which show the *z*-statistic maps (i.e., bootstrap test statistics) since the model “coefficients” obtained using the approach described by Zhang et al., (2014) are not true model weights, and in our experience the maps obtained using this approach for non-linear support vector models may contain very extreme values that make it difficult to visualize the overall pattern of weights. Plots shown to the right of the maps mirror those reported in the main text. For regression models, these show from left to right: the full-sample cross-validation correlation (as a scatterplot), the permutation test results for the in-sample model fit, the mean/SD of cross-validation R-squared values for each repeat of the cross-validation procedure, and the permutation test results for the cross-validation analysis. For classification models, these show from left to right: the full-sample cross-validation Fisher Exact Test (as a confusion matrix), the permutation test results for the in-sample model fit, the mean/SD of cross-validation area under ROC curves for each repeat of the cross-validation procedure, and the permutation test results for the cross-validation analysis.

**Supplemental Figure 1. Mass-Univariate Lesion-Symptom Mapping Results**

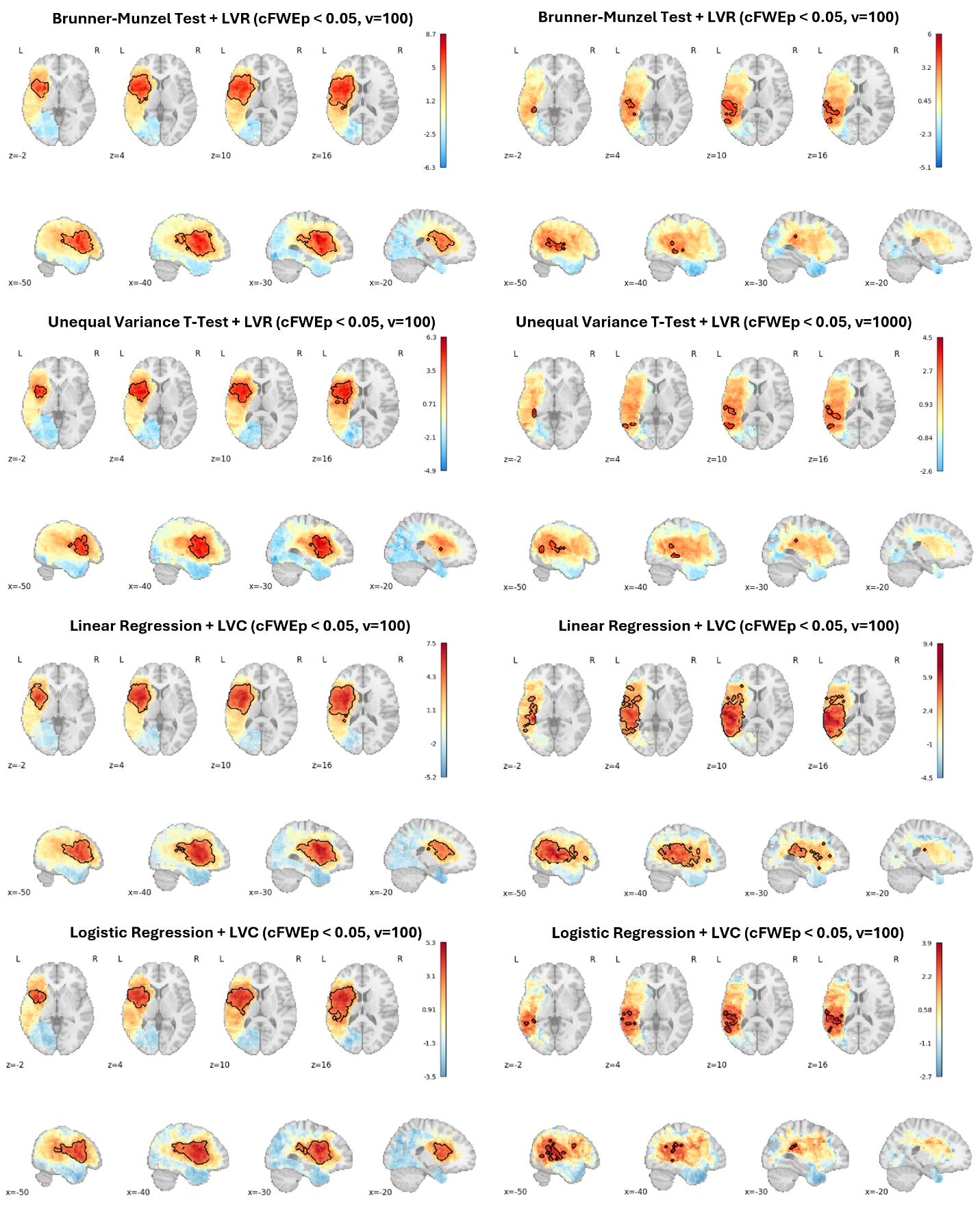

**Supplemental Figure 2. Multivariate Lesion Regression Results for Token Test (No DTLVC)**

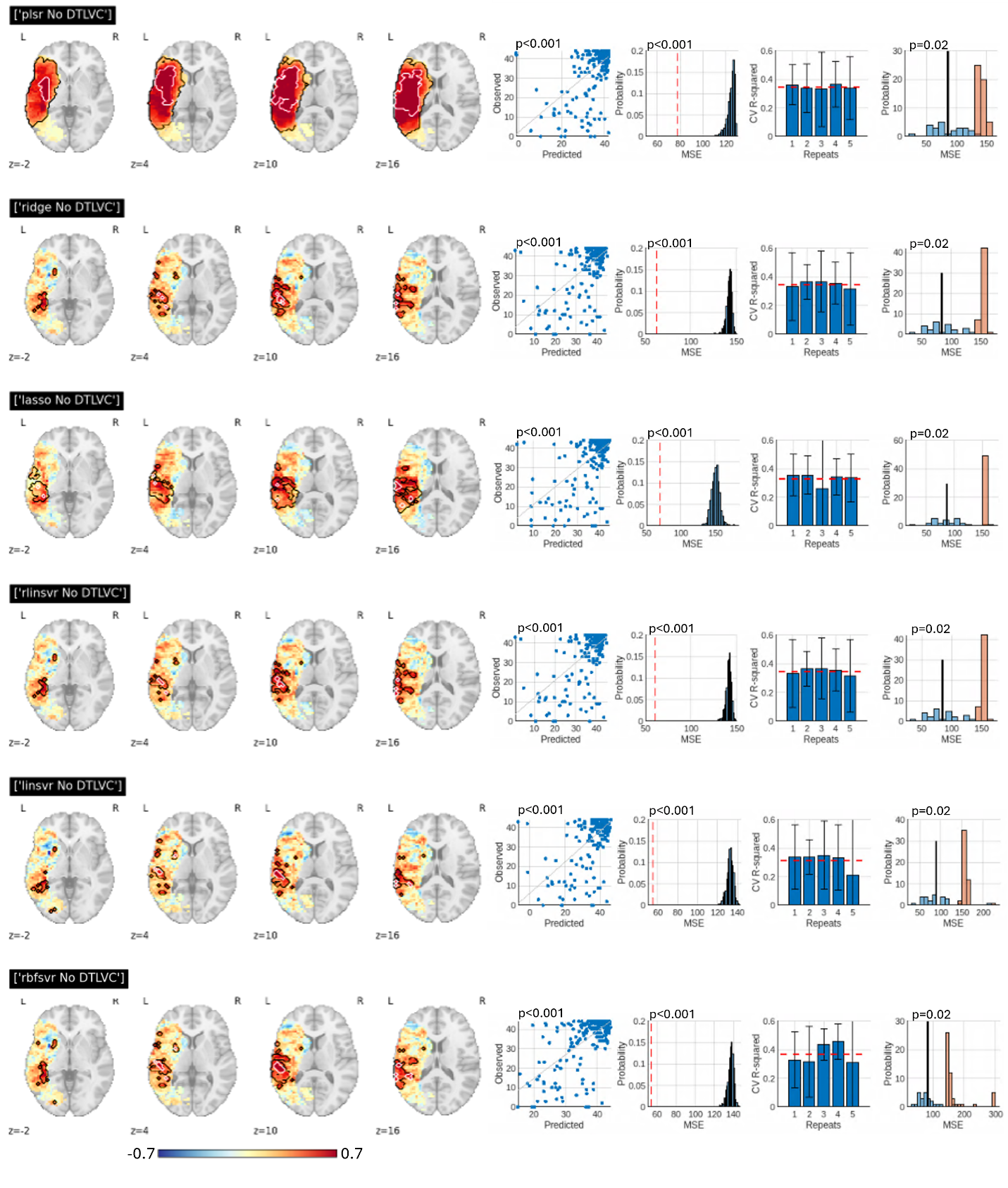

**Supplemental Figure 3. Multivariate Lesion Regression Results for Token Test (+ DTLVC)**

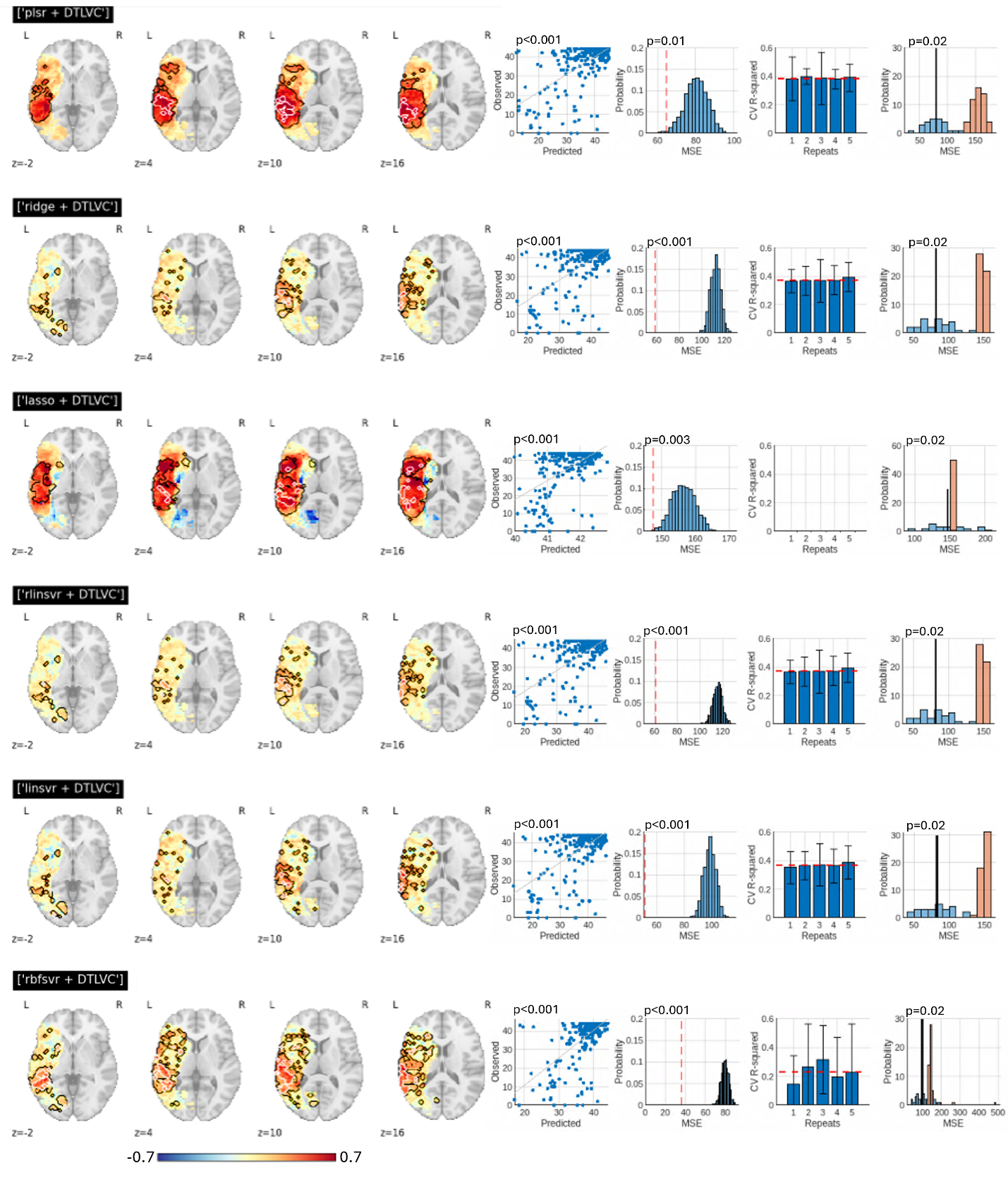

**Supplemental Figure 4. Multivariate Lesion Classification Results for Token Test (No DTLVC)**

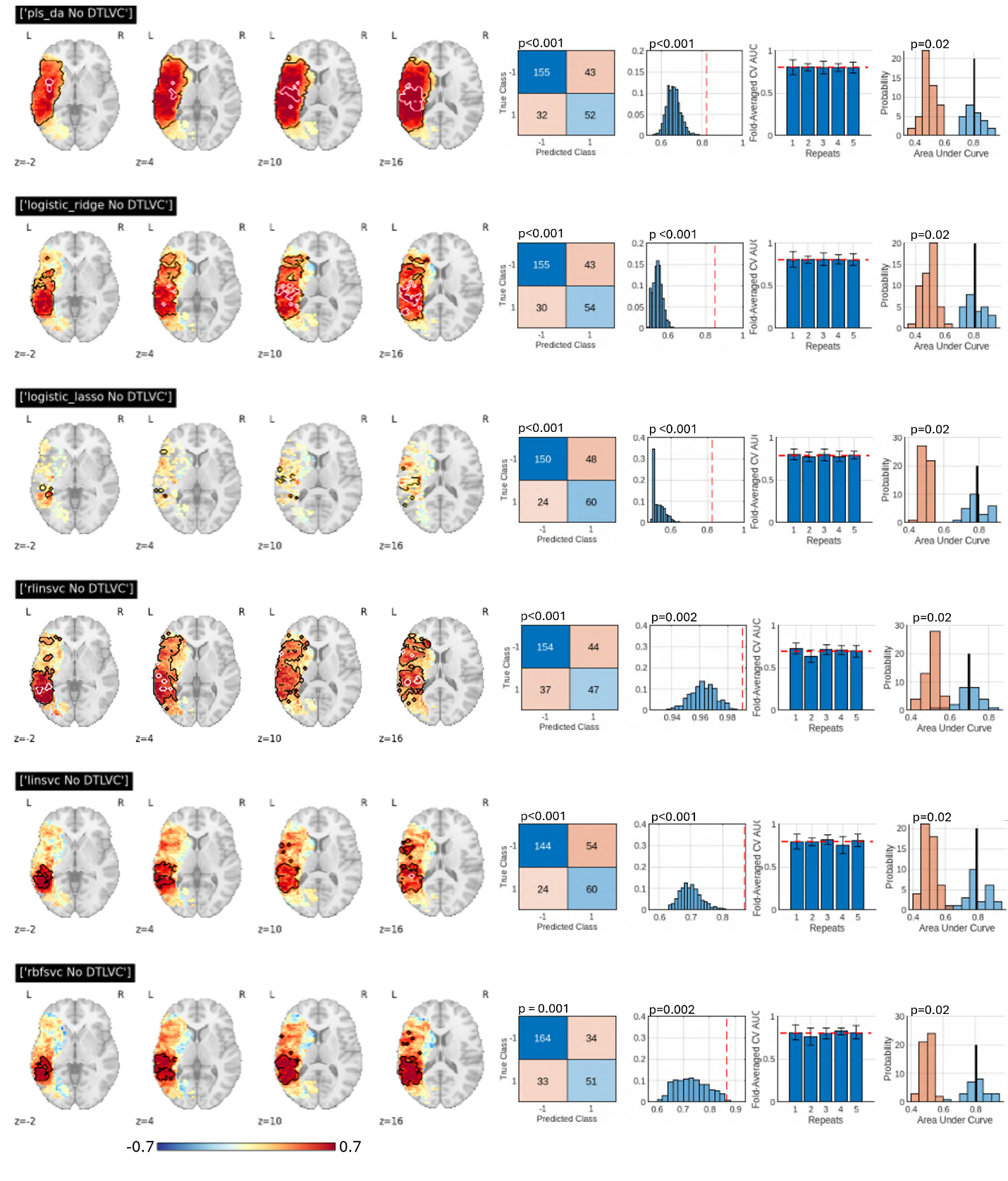

**Supplemental Figure 5. Multivariate Lesion Classification Results for Token Test (+ DTLVC)**

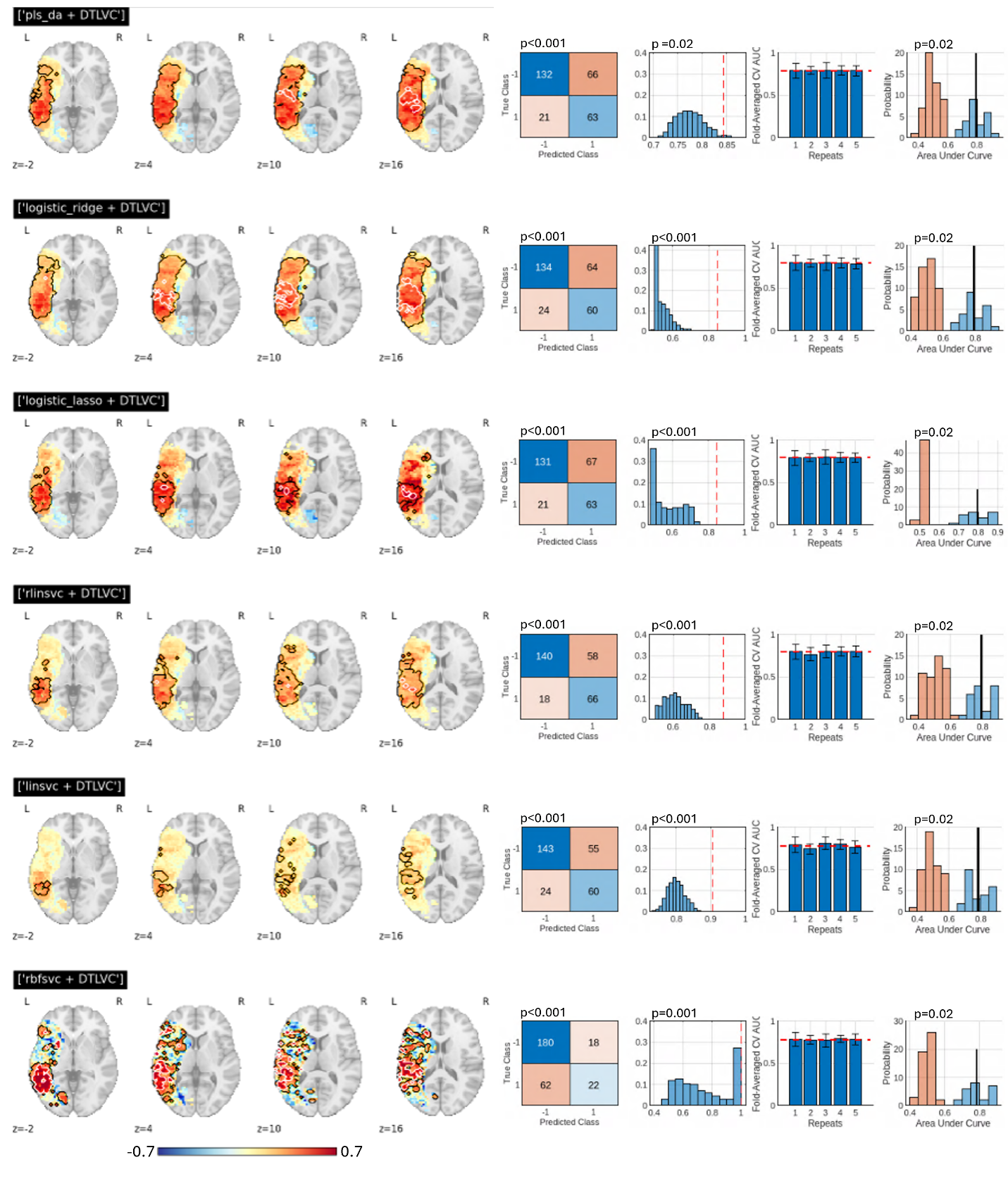

**Supplemental Figure 6. Multivariate Lesion Regression Results for COWA (No DTLVC)**

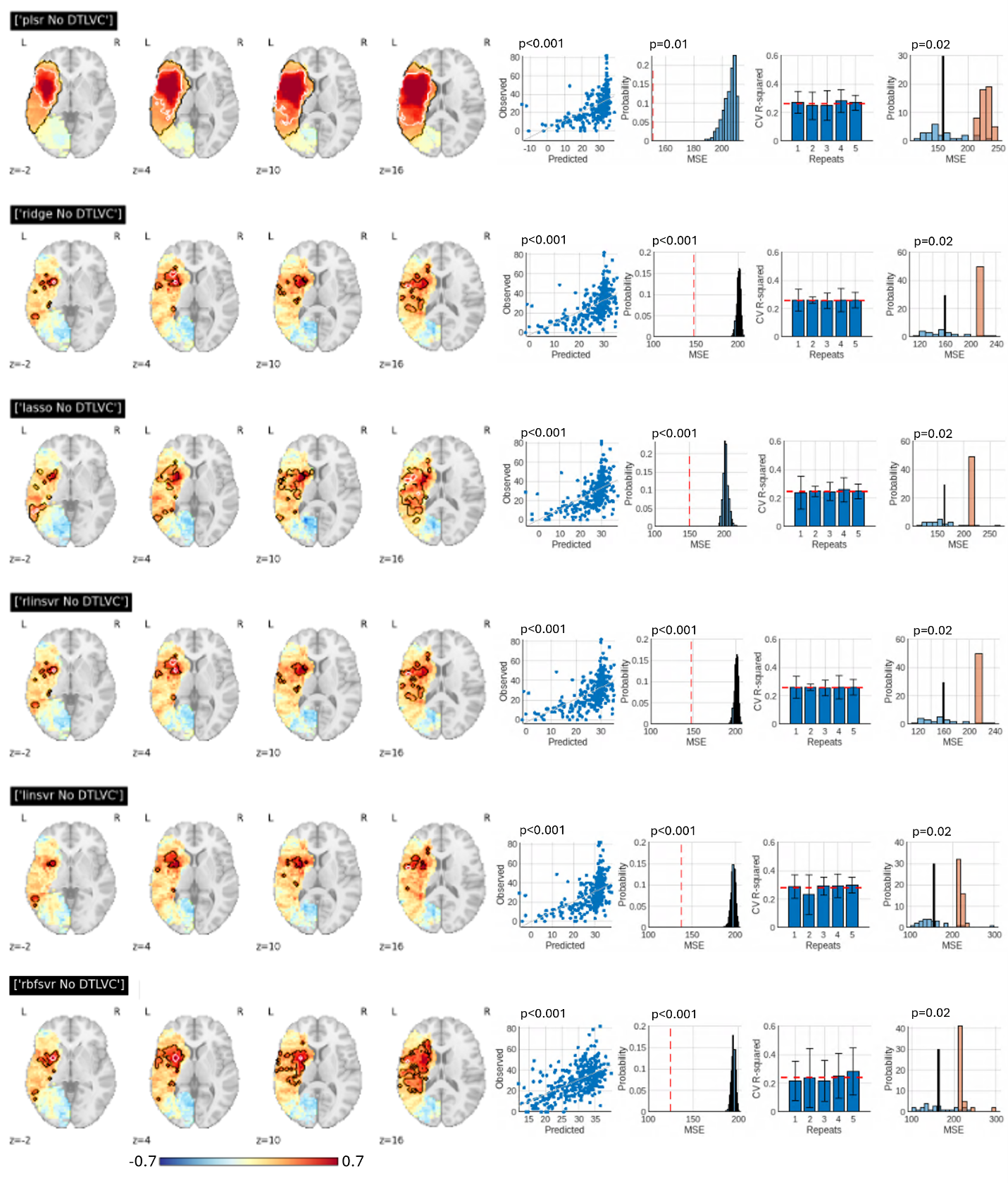

**Supplemental Figure 7. Multivariate Lesion Regression Results for COWA (+ DTLVC)**

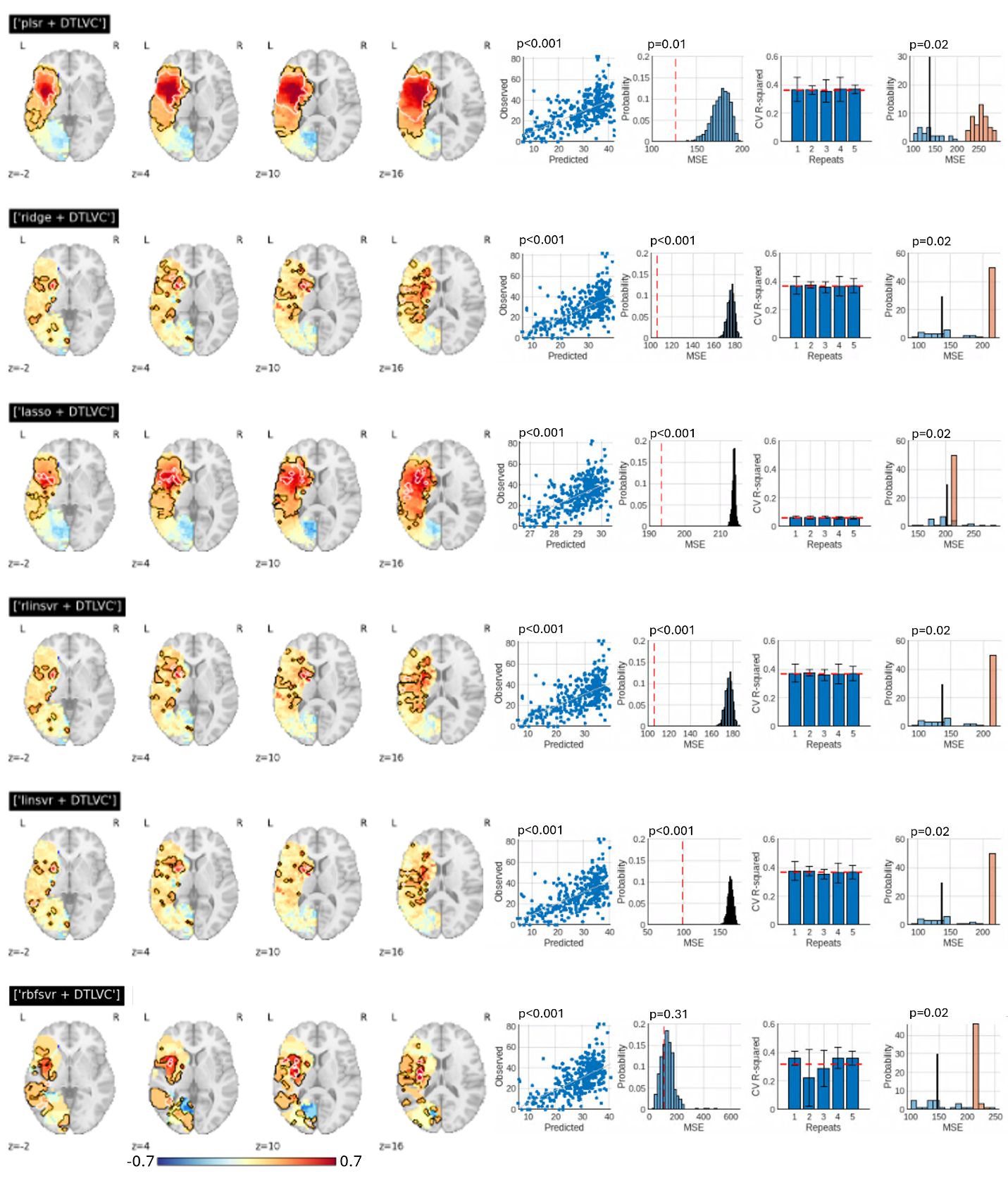

**Supplemental Figure 8. Multivariate Lesion Classification Results for COWA (No DTLVC)**

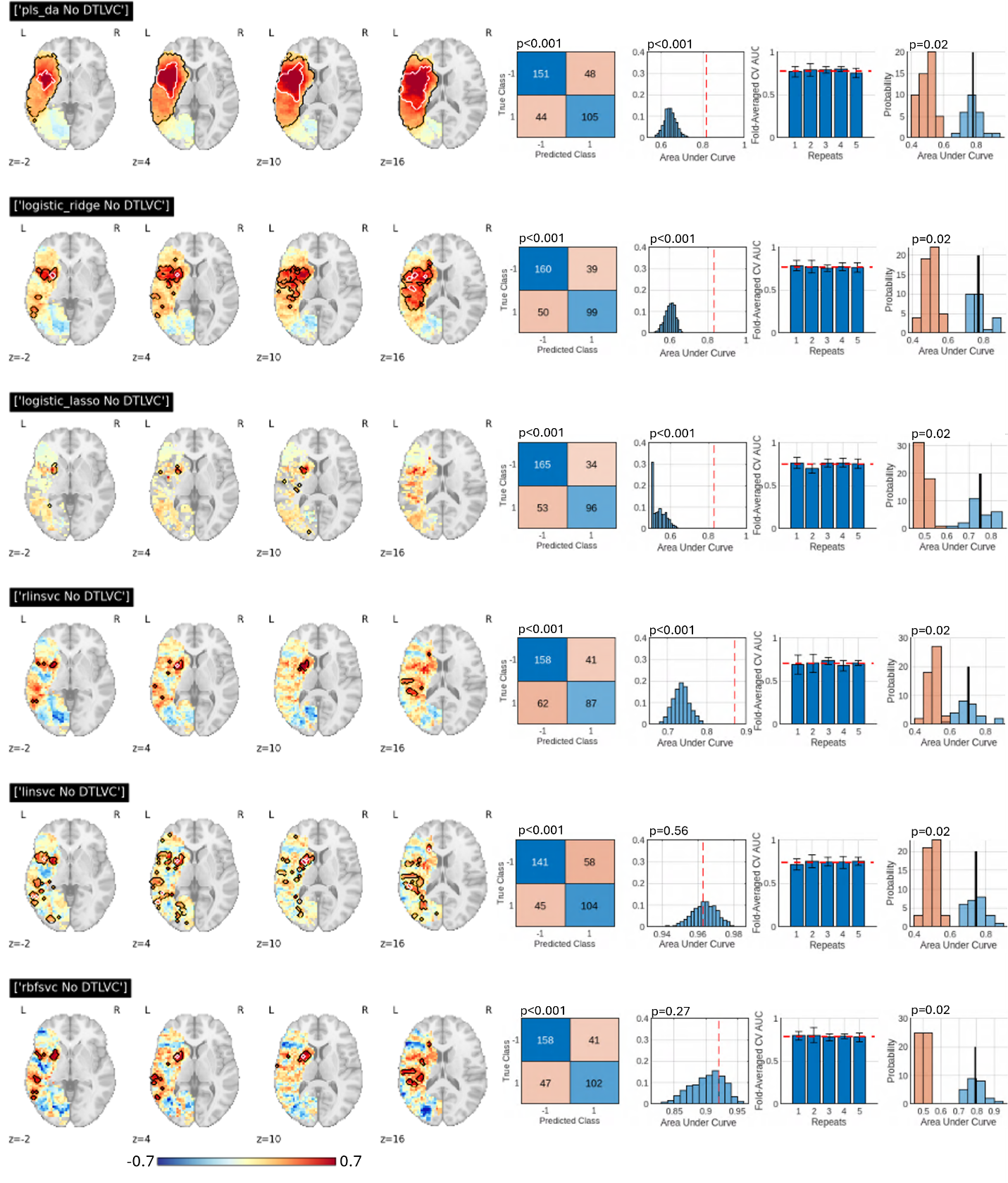

**Supplemental Figure 9. Multivariate Lesion Classification Results for COWA (+ DTLVC)**

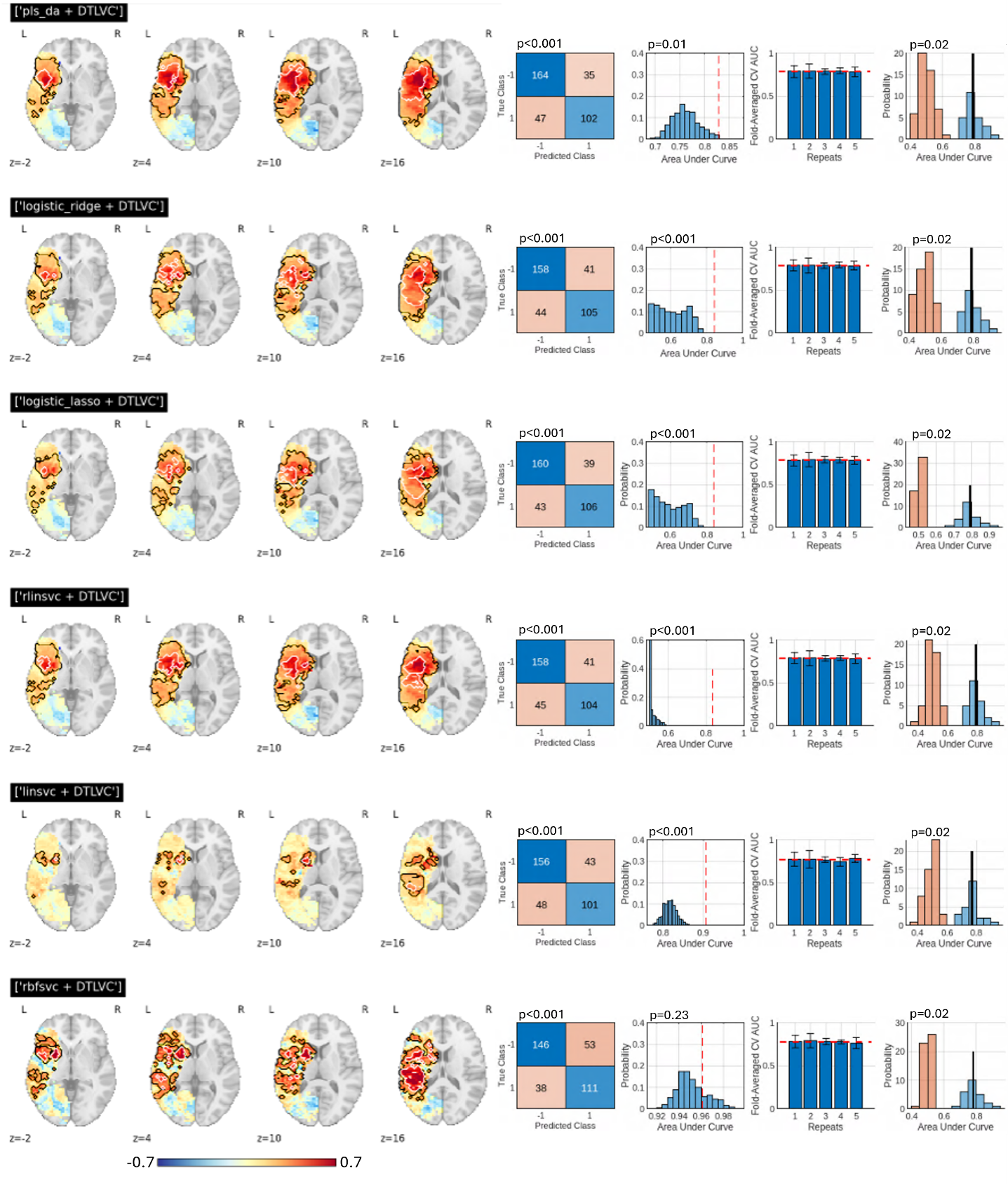
